# Microglia actively remodels adult hippocampal neurogenesis through the phagocytosis secretome

**DOI:** 10.1101/583849

**Authors:** Irune Diaz-Aparicio, Iñaki Paris, Virginia Sierra-Torre, Ainhoa Plaza-Zabala, Noelia Rodríguez-Iglesias, Mar Márquez-Ropero, Sol Beccari, Oihane Abiega, Elena Alberdi, Carlos Matute, Irantzu Bernales, Angela Schulz, Lilla Otrokocsi, Beata Sperlagh, Kaisa E. Happonen, Greg Lemke, Mirjana Maletic-Savatic, Jorge Valero, Amanda Sierra

**Author notes:** **Corresponding authors** Amanda Sierra, PhD, Research Professor, Achucarro Basque Center for Neuroscience, Science Park of the University of the Basque Country (UPV/EHU), Sede Building, 3^rd^ floor, Leioa, Bizkaia, 48940, Spain, Jorge Valero, PhD, Research Fellow, Achucarro Basque Center for Neuroscience, Science Park of the University of the Basque Country (UPV/EHU), Sede Building, 3^rd^ floor, Leioa, Bizkaia, 48940, Spain.

## Abstract

During adult hippocampal neurogenesis, the majority of newborn cells undergo apoptosis and are rapidly phagocytosed by resident microglia to prevent the spillover of their intracellular contents. Here, we propose that phagocytosis is not merely a passive process of corpse removal but has an active role in maintaining adult hippocampal neurogenesis. First, we found that neurogenesis was disrupted in mice chronically deficient for two microglial phagocytosis pathways (P2Y12 and MerTK/Axl), but was transiently increased in mice in which MerTK expression was conditionally downregulated. Next, we performed a transcriptomic analysis of microglial phagocytosis in vitro and identified genes involved in metabolism, chromatin remodeling, and neurogenesis-related functions. Finally, we discovered that the secretome of phagocytic microglia limits the production of new neurons both in vivo and in vitro. Our data suggest that reprogrammed phagocytic microglia act as a sensor of local cell death, modulating the balance between cell proliferation and cell survival in the neurogenic niche, thereby supporting the long-term maintenance of adult hippocampal neurogenesis.

**GRAPHICAL ABSTRACT:** 

## INTRODUCTION

Neurogenesis, or the formation of new neurons, is a complex process that extends throughout adulthood in specific regions of the mammalian brain. Here we focus on the subgranular zone (SGZ) of the hippocampus, which generates newborn granule cells in rodents (Ehninger and Kempermann, 2008) and humans (Moreno-Jimenez et al., 2019). Adult hippocampal neurogenesis comprises different steps through which radial neural stem cells (rNSCs) proliferate, differentiate into neuron-committed cells (neuroblasts) and migrate until they eventually integrate into the existing circuitry and gradually acquire physiological neuronal properties (Kempermann et al., 2004). Nowadays, newly generated neurons are strongly suggested to contribute to hippocampus-dependent learning and memory, among other functions (Deng et al., 2010).

Multiple endogenous factors regulate the proliferation, survival, differentiation and integration of the new neurons in the adult hippocampus. In the cellular niche, one key element is microglia, the resident macrophages of the nervous system that coordinate the brain inflammatory response. The detrimental effect of neuroinflammation on neurogenesis is well described, and is mediated by inflammatory cytokines such as interleukin 1 beta (IL-1β), tumor necrosis factor alpha (TNFα) and interleukin 6 (IL-6) (Ekdahl et al., 2003; Monje et al., 2003). Although the main function of inflammatory cytokines is to defend the organism, their excessive and chronic release leads to detrimental consequences, including induction of apoptosis in neurons and glial cells and the release of neurotoxic factors such as ROS (reactive oxygen species) and NO (nitric oxide) (Sastre et al., 2006; Smith et al., 2012), factors that could also contribute to the detrimental effects of inflammation on neurogenesis.

Microglia also beneficially affects neurogenesis, as they are capable of producing factors that modulate proliferation or survival of different cells within the neuronal lineage. In vitro studies demonstrate that cultured microglia promote differentiation of precursor cells (Aarum et al., 2003), while microglia conditioned media enhances neuroblast production and neurons survival (Morgan et al., 2004; Walton et al., 2006). Throughout the literature, a limited number of growth factors, such as FGF-2 (Fibroblast growth factor-2) or IGF-1 (Insulin-like growth factor-1), have been described to be produced by microglia under different conditions (Sierra et al., 2014). Furthermore, microglia were suggested to inhibit the proliferation of hippocampal rNSCs, as their number inversely correlates with adult hippocampal neurogenesis (Gebara et al., 2013). Recently, experiments using diphteria toxin-induced ablation of microglia propose that microglia are essential for neuroblast survival (Kreisel et al., 2018) but the mechanisms underlying the regulation of hippocampal neurogenesis by microglia are still unexplored both in vitro and especially in vivo (Sierra et al., 2014).

Here, we focus on another major role of microglia in the adult hippocampal neurogenic niche: the removal of the excess of newborn cells through phagocytosis (Sierra et al., 2010). Hippocampal newborn cells undergo apoptosis in the first few days of life through adulthood (Beccari et al., 2017) and are immediately recognized and degraded by ‘unchallenged’ microglia, in a process that lasts less than 1.5 hours (Sierra et al., 2010). Microglia are the brain professional phagocytes and their branches are highly motile, supporting a constant scanning of the brain searching for alterations in homeostasis (Sierra et al., 2013). In addition, microglia are equipped with receptors for the ‘find-me’ and ‘eat-me’ signals produced by apoptotic cells, which exert a positive chemotaxic response that results in the engulfment of the apoptotic cell in a phagosome that will subsequently fuse with lysosomes for its complete degradation (Sierra et al., 2013). Therefore, microglial phagocytosis is tightly coupled to apoptosis (Abiega et al., 2016) and prevents the release of toxic intracellular contents (Nagata et al., 2010) and thus, this process is essential to avoid alterations of the surrounding tissue.

Moreover, there is also increasing evidence that microglia are able to produce different trophic factors after apoptotic cell phagocytosis. Phagocytic microglia in culture are capable of producing transforming growth factor beta (TGF-β) as well as nerve growth factor (NGF) (De Simone et al., 2003). Similarly, hepatic macrophages produce vascular endothelial growth factor (VEGF) upon phagocytosis (Golpon et al., 2004). These three factors (TGF-β, NGF and VEGF) are potent regulators of hippocampal neurogenesis in vivo (Buckwalter et al., 2006; Cao et al., 2004; Kreisel et al., 2018). In this study, we propose that microglial phagocytosis does not conclude with the physical elimination of apoptotic cells, but is followed by a coordinated transcriptional program that triggers the production of neurogenic modulatory factors, which directly contribute to the maintenance and correct regulation of the adult hippocampal neurogenic cascade. Using constitutive and inducible knock-out (KO) mice to abolish different phagocytosis-related receptors, we discovered that chronic phagocytosis deficiency disrupts neurogenesis. In addition, using a combined in vitro and in vivo based experimental strategy, we found that the secretome of phagocytic microglia limits the production of new neurons in order to maintain the homeostasis of the adult hippocampal neurogenic niche.

## RESULTS

### Chronic impairment of microglial phagocytosis reduces adult hippocampal neurogenesis

To examine the impact of microglial phagocytosis on adult hippocampal neurogenesis in vivo, we focused on two signaling pathways involved on phagocytosis: P2Y12 (purinergic receptor type Y12, which recognizes ADP) and the TAM family tyrosine kinases MerTK and Axl (which bind to phosphatidylserine adapter/bridging molecules Growth arrest specific factor 6 (Gas6) and Protein S (Pros1))(Elliott et al., 2009; Fourgeaud et al., 2016). We used two transgenic mouse models in which these proteins are constitutively knocked out (KO: P2Y12 KO and MerTK/Axl KO). We decided to study the impact of phagocytosis in young mice (1m), as neurogenesis and apoptosis of newborn cells and subsequent phagocytosis by microglia rapidly declines with age (Beccari et al., 2017; Sierra et al., 2010). Apoptotic cells were defined as pyknotic/karryorhectic nuclei labeled with the DNA dye DAPI, which we have previously characterized to express other apoptosis markers such as activated caspase 3 and fractin (Sierra et al., 2010). First, we assessed phagocytosis in the hippocampus of the two KO models by quantifying the Ph index (the percentage of apoptotic cells engulfed by microglia), which is around 90% in physiological conditions (Abiega et al., 2016), and found significantly lower Ph index in the two KO models (74.8 ± 0.9% for P2Y12, 61.5 ± 1.6% for MerTK/Axl) (Figure 1A-C). In addition, the microglial Ph capacity (weighted average of the number of pouches containing apoptotic cells per microglia, i.e., the average number of phagocytic pouches per microglia) was significantly reduced in both KO models (Figure 1A-C). Nonetheless, the phagocytosis reduction was small, possibly due to compensatory mechanisms resulting from the chronic depletion, and we only detected the expected increase of apoptotic cells in MerTK/Axl KO mice **(Supplementary Figure 1A)**, possibly indicating not a complete dysfunction but a slowdown of phagocytosis.

**Figure 1.**
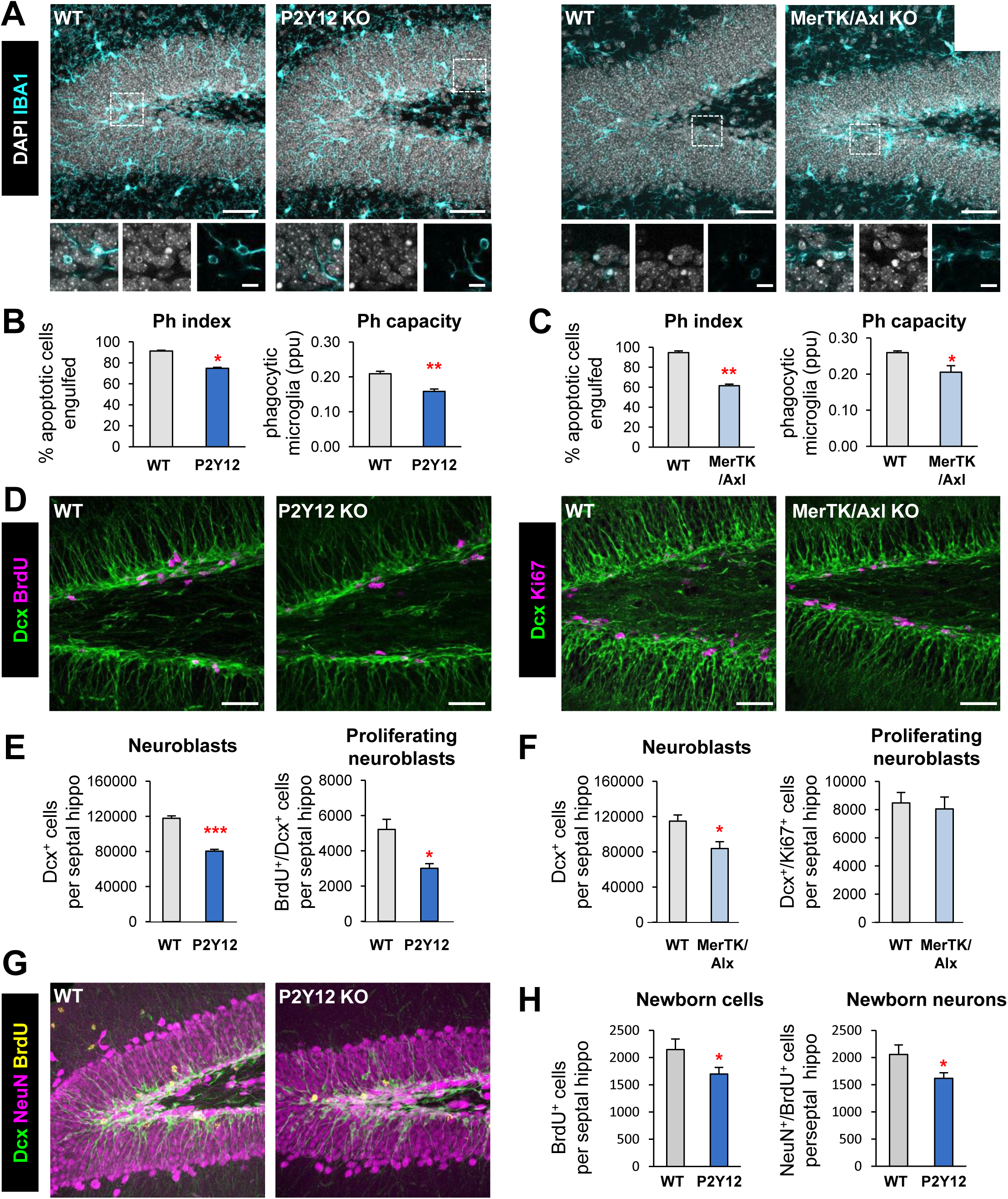
Chronic microglial phagocytosis impairment reduces adult hippocampal neurogenesis. **[A]** Representative maximum projection of confocal z-stack of P2Y12 and MerTK/Axl KO mice immunofluorescence in the mouse hippocampal DG at 1 month (1m). Microglia were labeled with Iba1 (cyan) and apoptotic nuclei were detected by pyknosis/karyorrhexis (white, DAPI). **[B, C]** Percentage of apoptotic cells engulfed (Ph index) and weighted average of the percentage of microglia with phagocytic pouches (Ph capacity) in P2Y12 KO mice (B) and MerTK/Axl KO mice (C). **[D]** Representative confocal z-stack of P2Y12 and MerTK/Axl KO mice immunofluorescence in the mouse hippocampal DG at 1m. Neuroblasts were labeled with DCX (green) and proliferation was labeled with either BrdU (150mg/kg, 24h) or Ki67 (magenta). **[E]** Neuroblast and neuroblast proliferation in 1m P2Y12 KO mice. **[F]** Neuroblast and neuroblast proliferation in 1m MerTK/Axl KO mice. **[G]** Representative confocal z-stack of P2Y12 KO mice immunofluorescence in the mouse hippocampal DG in physiological conditions at 2m. Neuroblasts were labeled with DCX (green), neurons were labeled with NeuN (magenta) and proliferation was labeled with BrdU (yellow, 4w after BrdU injection). **[H]** New cells and new neurons (NeuN^+^, BrdU^+^) in 2m P2Y12 KO mice, 4w after the BrdU injection. Scale bars, 50µm (A, D, G; inserts 10μm); z=20μm (A left panel), 17μm (A right panel), 7μm (D left panel), 10μm (D right panel), 20μm (G). N=3-4 mice (B, C, E, F), N=5 mice (H), Bars represent mean ± SEM. * indicates p < 0.05; ** indicates p < 0.01; *** indicates p < 0.001 by Student’s t-Test. Only significant effects are shown.

Next, we examined hippocampal neurogenesis in these phagocytosis impaired KO models and observed that all two showed a significant decrease in the population of neuroblasts, labeled with doublecortin (DCX), compared to wild type (WT) controls. In addition, we assessed proliferation by quantifying the number of dividing cells using either bromodeoxyuridine (BrdU) an analog of thymidine that gets incorporated into the DNA during S phase of dividing cells (mice were injected with 150mg/kg and sacrificed 24h later); or the proliferation marker Ki67^+^ (Scholzen and Gerdes, 2000). P2Y12 KO mice had a decrease in both neuroblasts (reduction of 31.7 ± 2.7%) and neuroblast proliferation (reduction of 39.3 ± 9.8%), and MerTK/Axl KO mice showed a reduction in neuroblasts (reduction of 26.2 ± 9.4%) compared to WT mice (Figure 1D-G). We further studied the formation of newborn neurons using BrdU pulse-and-chase in the most robust model, the P2Y12 KO. Four weeks after the BrdU injection, when the mice were 2m, both total BrdU^+^ cells and newborn neurons (NeuN^+^/BrdU^+^) were reduced in P2Y12 KO mice compared to WT mice (reduction of 24.6 ± 9.7%) (Figure 1G, H), in parallel to a decrease in phagocytosis (Ph index and Ph capacity; **Supplementary Figure 1B**).

To confirm the specificity of these results, we analyzed the expression of P2Y12, MerTK and Axl in FACS sorted cells from 1m fms-EGFP mice, in which microglia is labeled with EGFP. We found that, P2Y12 and MerTK, but not Axl, were highly expressed in microglia compared to other cells of the brain parenchyma **(Supplementary Figure 1C, D)**, suggesting that the disruption of neurogenesis in the KO models might be attributable to the lack of these receptors in microglia. While these receptors regulate multiple features of microglial physiology (Elliott et al., 2009; Fourgeaud et al., 2016), their similar neurogenesis impairment suggests that an intact microglial phagocytosis is necessary for the long-term maintenance of hippocampal neurogenesis.

### Acute microglial phagocytosis impairment transiently increases adult hippocampal neurogenesis

We then studied the effect of acute phagocytosis blockage on neurogenesis using an inducible MerTK KO model (generated by crossing *Mertk*^*fl/fl*^ to *Cx3cr1*^*CreER*/+^ mice) (Fourgeaud et al., 2016). Mice received tamoxifen (two 75mg/kg ip injections) at postnatal days p21 and p23 to induce microglial-specific Cre-mediated depletion of *Mertk*, and one injection of BrdU at p28 to label dividing cells. Phagocytosis and neurogenesis were analyzed at 1d and 4 weeks after BrdU administration (i.e., when mice were 1m and 2m, respectively; Figure 2A, B). Phagocytosis was strongly reduced in the inducible KO mice (iKO), as shown by the Ph index (73.0% ± 3.7 and 72.9% ± 2.6% reduction at 1d and 4w, respectively) and the Ph capacity (54.5% ± 3.1% and 65.6% ± 11.0% reduction at 1d and 4w, respectively), together with an accumulation of apoptotic cells (Figure 2C-F, **Supplementary Figure 1E**). Concomitantly, 1d after the BrdU injection the number of proliferating BrdU^+^ cells and the number of DCX^+^, BrdU^+^, proliferating neuroblasts (51.8% ± 10.2% and 55.4% ± 12.2% increase, respectively), while there was no change in the total number of neuroblasts in MerTK iKO compared to WT mice (Figure 2G-I). However, at 4w after BrdU injection there were no significant changes in the total number of newborn cells nor newborn neurons (BrdU^+^, NeuN^+^) (Figure 2J, K). These data suggested that the excess of BrdU cells formed at 1d were lost at 4w, and indeed the net yield of newborn cells was significantly lower in iKO compared to WT mice (43.1% ± 4.5% reduction; Figure 2L). The transient increase in early neuroblasts after acute impairment of phagocytosis in the iKO model, together with the chronic reduction of neurogenesis in the constitutive KO models, suggest that the microglial phagocytosis of newborn cells participates in a feedback loop that maintains the homeostasis of adult hippocampal neurogenesis.

**Figure 2.**
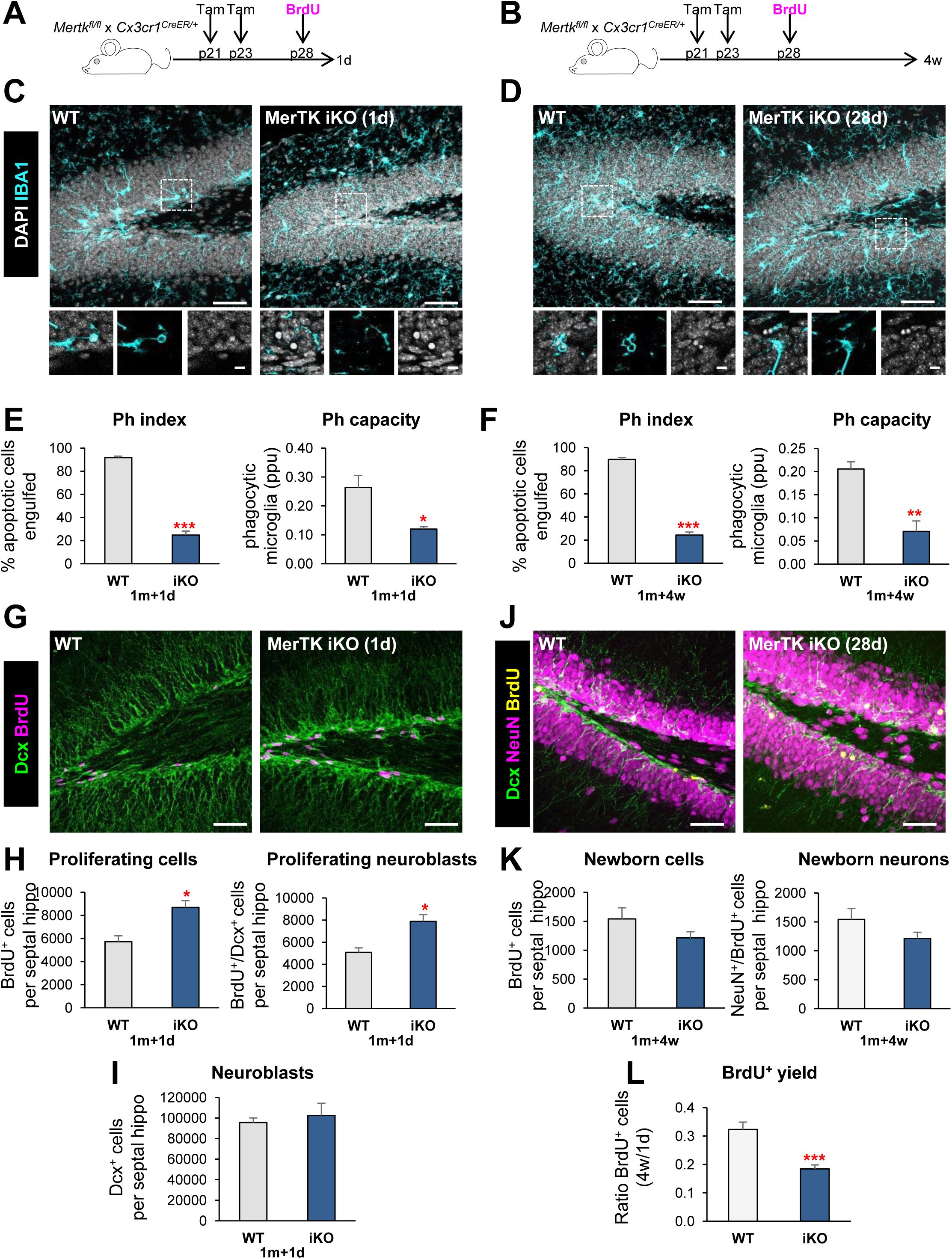
Acute microglial phagocytosis impairment transiently increases adult hippocampal neurogenesis. **[A, B]** Experimental design in microglial-specific MerTK inducible KO mice (iKO), generated by crossing *Mertk*^*fl/fl*^ and *Cx3cr1*^*CreER*^ mice treated with tamoxifen at p21 and p23 and injected with BrdU at p28 (100mg/kg). Mice were sacrificed at 1d (A) or 4w (B) after the BrdU injection. **[C, D]** Representative maximum projection of confocal z-stack of MerTK iKO mice immunofluorescence in the mouse hippocampal DG at 1d (C) and 4w (D). Microglia were labeled with Iba1 (cyan) and apoptotic nuclei were detected by pyknosis/karyorrhexis (white, DAPI). **[E, F]** Percentage of apoptotic cells engulfed (Ph index) and weighted average of the percentage of microglia with phagocytic pouches (Ph capacity) in MerTK iKO mice at 1d (E) and 4w (F). **[G]** Representative confocal z-stack of MerTK iKO mice immunofluorescence in the mouse hippocampal DG at 1d. Neuroblasts were labeled with DCX (green) and proliferation was detected with BrdU (magenta). **[H]** Newborn cells (BrdU^+^) and newborn neuroblasts (DCX^+^,BrdU^+^) in MerTK iKO mice at 1d post-BrdU. **[I]** Neuroblasts (DCX^+^) in MerTK iKO mice at 1d post-BrdU. **[J]** Representative confocal z-stack of MerTK iKO mice immunofluorescence in the mouse hippocampal DG at 4w post-BrdU. Neuroblasts were labeled with DCX (green), neurons were labeled with NeuN (magenta) and proliferation was labeled with BrdU (yellow). **[K]** Newborn cells (BrdU^+^) and newborn neurons (NeuN^+^,BrdU^+^) in MerTK iKO mice at 4w post-BrdU. **[L]** BrdU+ yield was calculated as a ratio of the BrdU^+^ cells at 4w over the average BrdU^+^ cells of each group at 1d after injection. Scale bars, 50µm (C, D, G, J; inserts 10μm); z=16.1μm (C, D), z=7mm (G), z=23.1μm (J). N=3 mice (E, H, I), N=4-6 mice (F, K, L). Bars represent mean ± SEM. * indicates p < 0.05; ** indicates p < 0.01; *** indicates p < 0.001 by Student’s t-Test. Only significant effects are shown.

### Phagocytosis of apoptotic cells triggers the expression of neurogenic modulatory factors by microglia in vitro

To study the mechanism by which microglial phagocytosis maintains neurogenesis in the long term, we developed a xenogeneic in vitro model of phagocytosis of apoptotic cells (Beccari et al., 2018), in which mouse primary microglia were fed for different lengths of time (1-24h) with a human neuronal line (SH-SY5Y), previously labeled with CM-DiI and treated with staurosporine (STP; 4h, 3µM) to induce apoptosis (Figure 3A-C). When fed with apoptotic human cells, microglia were phagocytic as early as 1h (30.2% ± 5.8%), a percentage that kept increasing until 24h (83.8% ± 3.6%) (Figure 3C). We then performed a mouse-specific genome-wide transcriptomic analysis using gene expression mouse-specific arrays to compare naïve vs. phagocytic microglia. To disregard the possible detection of residual mRNA from the (human) apoptotic cells, we checked the sequence of the 60,000 array probes against the human transcriptome by BLAST. We found that 96% of the probes had low homology to the human transcriptome (MegaBlast homology < 5%). In addition, we analyzed the RNA integrity in apoptotic cells using a Bioanalyzer, and found that while naïve and phagocytic (24h) microglia had the expected 18S and 28S rRNAs profile, apoptotic SH-SY5Y cells (24h) showed a smear typical of RNA degradation **(Supplementary Figure 2A)**. Finally, we analyzed whether apoptotic cells could synthesize new mRNA using 5’-Fluorouridine (FU, a uridine analog that integrates at transcription sites) **(Supplementary Figure 2B)**. Apoptotic SH-SY5Y, unlike live cells, did not exhibit nuclear FU labeling, evidencing that they were not transcriptionally active. Altogether, the RNA profiling and analysis of transcription in apoptotic cells strongly suggests that although the gene arrays used could virtually detect up to 4% of mRNAs from human apoptotic cells, their lack of residual RNA would result solely in the detection of microglial-specific transcriptional changes after phagocytosis. In addition, the arrays findings were later validated by RT-qPCR using mouse-specific primers (see below).

**Figure 3.**
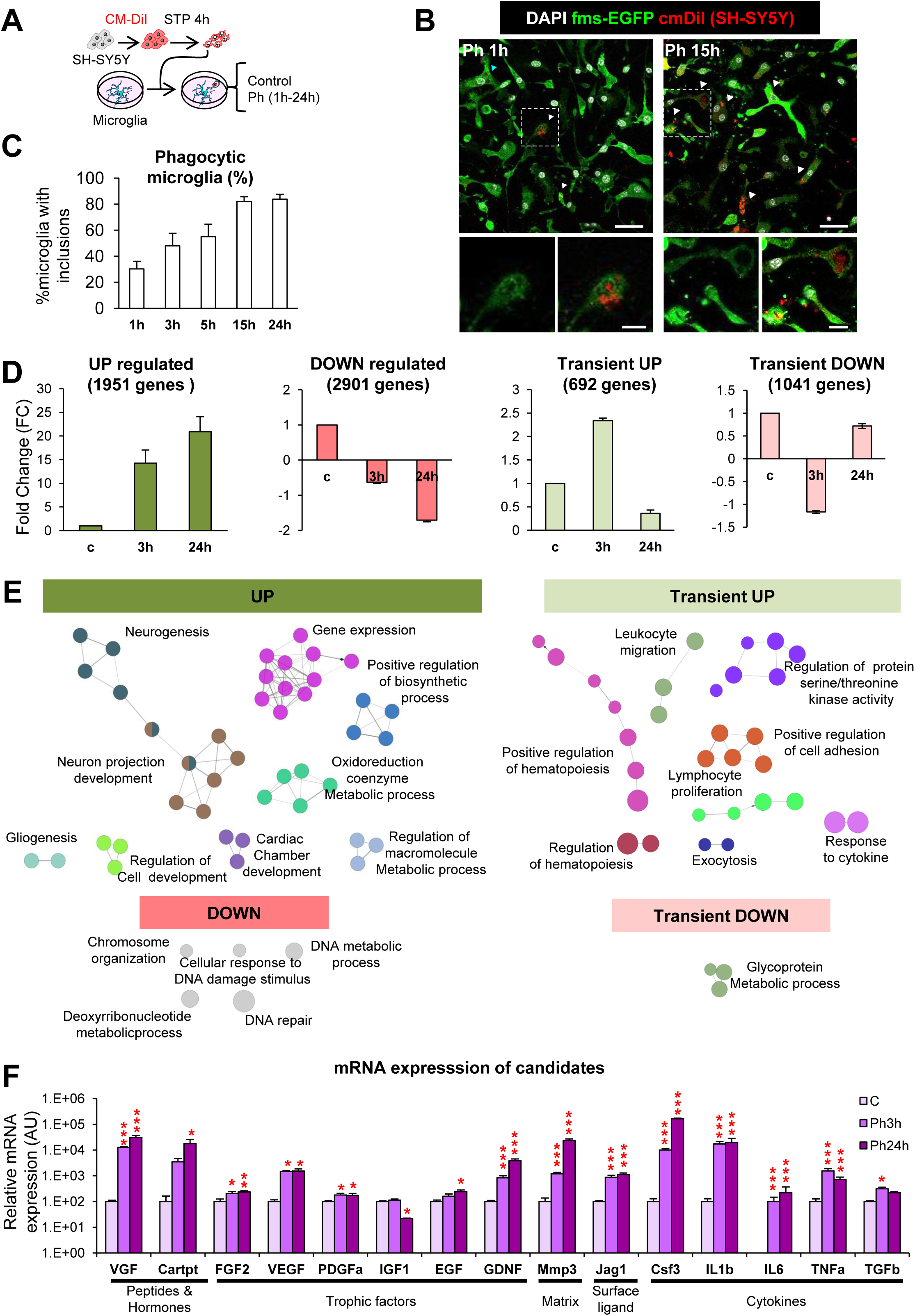
Phagocytosis assay with a human neural cell line (SH-SY5Y). **[A]** Experimental design of the phagocytosis assay. **[B]** Representative confocal microscopy images of primary microglia (GFP, green) fed with SH-SY5Y, which were previously labeled with CM-DiI (red) and treated with STP (4h, 3μm) for the induction of apoptosis (pyknosis/karyorrhexis, DAPI, white). N, microglial nucleus; arrowheads, phagocytosed apoptotic SH-SY5Y cells. **[C]** Percentage of microglia with CM-DiI and/or DAPI inclusions along a time course. Only fully closed pouches with particles within were identified as phagocytosis. **[D]** FC mean of the genes classified under the UP, DOWN, Transient UP and Transient DOWN regulation patterns. **[E]** Functional analysis of phagocytic microglia using ClueGO. Charts show the interactions among the significantly different functions for the four main expression patterns. Biological functions are visualized as colored nodes linked to related groups based on their kappa score level. The node size reflects the enrichment significance of the term and functionally related groups are linked. Non-grouped terms are shown in grey. **[F]** mRNA expression levels of the candidates selected for validation by RT-qPCR. N=4 independent experiments. HPRT was selected as a reference gene. Scale bars, 30µm (B; inserts, 10μm); z=6.3 µm. N=3 independent experiments (C, D, E, F), N=4 independent experiments (F). Bars represent mean ± SEM. * indicates p < 0.05; ** indicates p < 0.01; *** indicates p < 0.001 by Holm-Sidak posthoc test of (after one-way ANOVA was significant at p < 0.05). Only significant effects are shown.

Hence, we compared the genome-wide transcriptome naïve vs. phagocytic microglia (3h and 24h) using gene expression arrays **(Supplementary Figure 2C-E)**. Hierarchical Clustering (HCL) and Principal Component Analysis (PCA) of the transcriptome of control, Ph3h and Ph24h microglia showed strong differences in the expression clustering of the expression profile of three groups. To analyze which particular genes were different among the three experimental groups, we searched for array probes with significant changes over time using a p-corrected Benjamini-Hochberg value and a polynomial regression model to identify time patterns (Conesa et al., 2006). We obtained 10,000 significantly regulated probes with a restrictive criterion of R squared (rsq) > 0.9 and a fold change (FC) higher than 1.5 or lower than −1.5, eventually obtaining 6,585 genes that presented significant changes over time **(Supplementary Figure 2F)**.

We classified these genes according to four main expression patterns (Figure 3D): UP (up-regulation both at 3h and 24h), DOWN (down-regulation in both time points), Transient UP (up-regulation at 3h and down-regulation at 24h), and Transient DOWN (down-regulation at 3h and up-regulation at 24h). Genes in the UP expression pattern showed the largest average FC changes among all patterns, some of them even reaching 8,000FC at 24h. The rest of the patterns had on average more modest changes.

Next, we performed a functional analysis of the phagocytic microglia transcriptome using the ClueGO network (Figure 3E) and DAVID **(Supplementary Figure 3A)**. These analyses revealed a number of functional biological pathways associated with each of the four main expression patterns of phagocytic microglia including downregulation in pathways related to DNA and chromosomes and upregulation of different functions associated with metabolism and chromatin remodeling. Interestingly, different studies suggesting metabolic changes in phagocytes upon the uptake of apoptotic cells have recently emerged (Morioka et al., 2018). Importantly, for many upregulated genes, ClueGO revealed specific terms like ‘generation of neurons’, ‘neuron differentiation’ or ‘neuron development’ that were grouped under the term ‘neurogenesis’ and had a direct interrelation with ‘neuron projection development’ group.

We then focused on the identity of the neurogenesis-related genes using the following strategy **(Supplementary Figure 3B)**. To identify phagocytosis-related potential regulators of neurogenesis, we used MANGO (The Mammalian Adult Neurogenesis Gene Ontology), a database of 259 genes already described to be involved in the regulation of adult hippocampal neurogenesis (Overall et al., 2012). Out of the MANGO genes, 213 were found significantly regulated in our arrays. We then filtered those MANGO genes by disregarding those encoding for autologous proteins (i.e., acting on the same cell, such as transcription factors) and focusing on those encoding heterologous proteins (i.e., acting on neighbor cells, such as secreted molecules). We first applied this criterion to MANGO, and found 26 heterologous genes that had been previously identified to regulate adult hippocampal neurogenesis. To further extend the list of heterologous genes outside MANGO that could be potential regulators of neurogenesis, we looked into the filtered gene array list (rsq>0.9 and −1.5>FC>1.5), searched the GO terms associated with each MANGO heterologous gene, and selected those terms that could be related to different steps of the neurogenic process (proliferation, differentiation, migration, chemotaxis, survival and development). MANGO heterologous genes encompassed 57 different neurogenesis-related GO terms, such as growth factor activity (GO:0008083), nervous system development (GO:0007399), learning (GO:0007612), memory (GO:0007613), cell proliferation (GO:0008283), cell differentiation (GO:0030154), and neuron development (GO:0048666).

We then searched for heterologous genes belonging to each of these 57 categories in our arrays, using a less-restrictive list of 20,800 probes (rsq>0.7, no screening of FC). We finally obtained 224 genes with differential expression between naïve and phagocytic microglia, which were heterologous and whose function had been previously involved in neurogenesis (based on the GO terms). The 224 candidate genes were classified according to their main regulatory expression patterns: 94 UP, 73 DOWN, 28 Transient-UP and 29 Transient-DOWN genes **(Supplementary Figure 3B)**. In the four regulation patterns, the majority of the genes were categorized as trophic factors (between 23-29% in all regulatory patterns; **Supplementary Figure 4A, B)**. We also found cytokines, chemokines, peptides, and hormones as the main gene types of the candidates. Despite the fact that trophic factors were the largest percentage in each regulation pattern, they showed a similar and rather low mean FC compared to the other categories. Only the up-regulated peptides and hormones revealed a large mean of 800 FC at both 3h and 24h of phagocytosis. These data suggest that peptides and hormones were the most likely molecules to perform modulatory functions in the neurogenic niche by phagocytic microglia.

**Figure 4.**
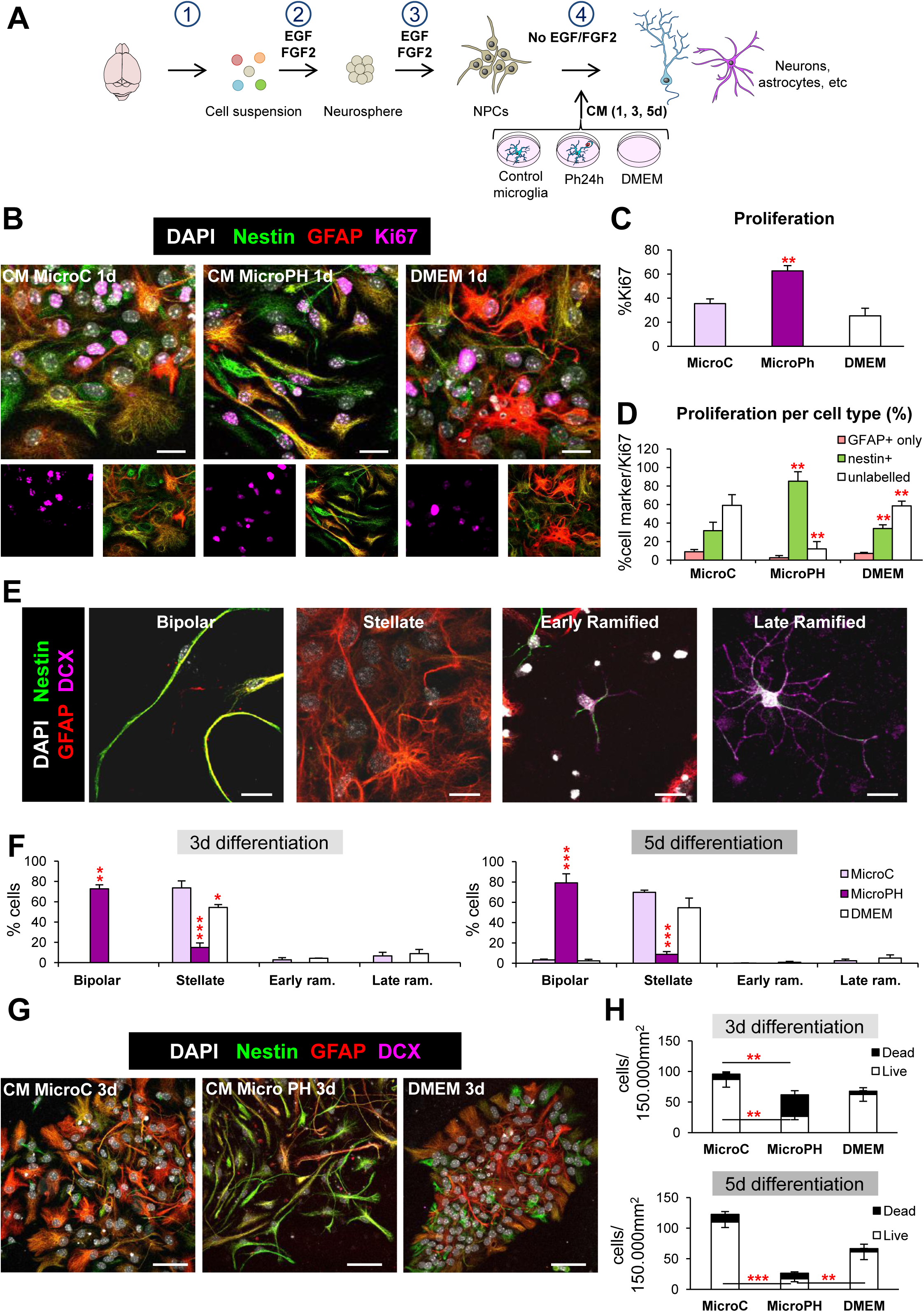
Effect of phagocytic microglia secreted factors on neurogenesis in vitro. **[A]** Experimental design of the in vitro neurogenesis assay: 1, brain disaggregation; 2, neurosphere proliferation; 3, dissociation, plating and proliferation of neuroprogenitors (NPCs) for 48h; 4, differentiation in the presence of conditioned media (CM) from control microglia (microC) or 24h phagocytic microglia (microPh). **[B]** Representative confocal microscopy images of NPCs treated with CM microC or microPH after 1d. DMEM was used as control. **[C]** Percentage of cells labeled with Ki67 over total cells labeled with DAPI. **[D]** Percentage of different cell markers over total Ki67 population: nestin^+^ (with or without GFAP), GFAP only, or unlabelled. **[E]** Representative confocal microscopy of the different morphologies observed in the neurogenesis assay images. **[F]** Percentage of the different cell types found after 3d and 5d treatment with CM microC or microPH. Early/late ram. refers to early/late ramified cells. **[G]** Representative confocal microscopy images of NPCs treated with CM microC or microPH after 3d. **[H]** Density of live and dead cells (determined by pyknosis/karyorrhexis) after CM treatment for 3d and 5d. Scale bars, 50µm (B), 20µm (E, G), z=11.9μm (B), 9µm (E, G). N= 3 independent experiments (C, D, F, H). Bars represent mean ± SEM. * indicates p < 0.05; ** indicates p < 0.01; *** indicates p < 0.001 by Holm-Sidak posthoc test vs MicroC group (after one-way ANOVA was significant at p < 0.05).

Finally, we validated the mRNA expression of the candidates in naïve and phagocytic microglia by RT-qPCR. We selected a subset of genes for validation based both on their high FC in the array and/or the well-known neurogenesis modulatory potential described in the literature. We found that the expression pattern of the selected candidates determined by RTqPCR was largely in agreement with that obtained in the arrays. Among the genes with the largest mRNA expression were the neuropeptide VGF (no abbreviation), the matrix metalloprotease 3 (MMP3), and the cytokine colony stimulating factor 3 (CSF3) (Figure 3F), reinforcing the notion that phagocytosis promotes the production of neurogenic modulators by microglia. Importantly, we delved into the requirements for triggering the neurogenic program: whether the program started in microglia as soon as they sensed the ‘find-me’ signals released from apoptotic cells or whether physical contact between microglia and apoptotic cells (tethering and engulfment) was necessary. To tease out these two possible mechanisms, we added conditioned media from apoptotic cells to naïve microglia and analyzed their mRNA expression of candidates 24h thereafter **(Supplementary Figure 2G)**. We found that candidates from extracellular space (matrix and cell-surface) and cytokines were highly up-regulated after culturing microglia with apoptotic cell media. The observed mRNA levels in this group were very similar to those at 24h of phagocytosis (Ph24h). However, the majority of ‘peptides and hormones’ and ‘trophic factors’ were not significantly changed. These results suggest that the neurogenic program triggered by microglial phagocytosis might be initiated by the detection of ‘find-me’ signals, but that further steps in the phagocytic process are required to develop the complete neurogenic program.

### The secretome from phagocytic and naïve microglia drives neuroprogenitor cells towards different fates in vitro

83.5% of the 224 heterologous candidates belonged to the secretome, suggesting an important role of the phagocytic microglial secretome on the modulation of neurogenesis. We thus directly tested the effect of the phagocytic microglial secretome on neurogenesis in vitro. To model neurogenesis, we used a monolayer of neuroprogenitor cell (NPC) cultures derived from disaggregated neurospheres, obtained from whole P0-P1 brains and allowed them to proliferate 48h in DMEM/F12 with trophic factors EGF/FGF2 (Babu et al., 2011). We first performed a neurogenesis differentiation assay in which NPCs were allowed to differentiate during a time course (1-5 days, d) in the presence of conditioned media from control (naïve microglia; CM microC) and phagocytic (24h phagocytic microglia, CM microPH) microglia (Figure 4A). DMEM was used as an internal control, as microglia were cultured in this media. After 1d of differentiation, cultures in CM microPH maintained higher levels of proliferation than CM microC as observed by a higher proportion of cells labeled with the proliferation marker Ki67^+^ (Figure 4B, C). Most of the proliferating cells in microPH cultures expressed nestin, a marker of progenitor cells, stem cells and reactive astrocytes (Encinas and Sierra, 2012; Lopez-Atalaya et al., 2018) (Figure 4D). We then followed the progeny of those populations at 3d and 5d and labeled them with cell identity markers: nestin, GFAP (Glial Fibrillary Acidic Protein), a marker of astrocytes (Encinas and Sierra, 2012) and DCX, a marker of neuroblasts (Brown et al., 2003) (Figure 4E). We found that CM microC treatment mainly produced GFAP^high^, nestin^+/−^, stellate cells both at 3d and 5d, as well as a small percentage of DCX ramified cells. In contrast, in CM microPH cultures the majority of the cells were nestin^high^, GFAP^+^, with a bipolar morphology (Figure 4F, G). Cell death (apoptosis) was observed in all conditions, as has been noted before in this type of cultures upon growth factor withdrawal-induced differentiation (Babu et al., 2011). However, higher rates of apoptosis were found in NPCs cultured in CM microPH, which resulted in a lower cell density (Figure 4H). Importantly, the majority (57%) of cell death-related upregulated heterologous genes were anti-apoptotic **(Table 1)**, suggesting that the phagocytic microglia secretome did not directly induce NPCs apoptosis. Taken together, the above results indicate that naïve microglia led to the production of stellate cells, resembling astrocytes in culture, and a small proportion of ramified DCX^+^ expressing cells, corresponding to an immature stage of the neuronal lineage. In contrast, phagocytic microglia drove NPCs towards a unique bipolar cell type, expressing nestin and GFAP but never DCX, which is characteristic of both astrocytes and undifferentiated progenitor cells (Encinas and Sierra, 2012).

### The secretome from phagocytic microglia drives neuroprogenitor cells towards an astrocytic phenotype in vitro

To precisely identify bipolar cells produced by CM microPH, we performed several studies to characterize their phenotype: labeling with the mature astrocytic marker (S100β) (Raponi et al., 2007), a multipotency assay, response to stimuli by calcium imaging, and Western blot analysis of neural and astrocyte-committed transcription factors.

First, we tested whether bipolar cells could be mature astrocytes and stained the CM-treated cultures with S100β, a marker of mature astrocytes and oligodendrocytes (Wang and Bordey, 2008) (Figure 5A). In CM microC few stellate cells were S100β^+^ and exhibited rather dim labeling, suggesting that they were still immature astrocytes (Figure 5B-D). In addition, in CM microC cultures, we observed a small percentage of potential oligodendrocytes, identified as cells that only expressed high levels of S100β and had several branched processes (Wang and Bordey, 2008). On the other hand, in CM microPH cultures oligodendrocyte-like cells were not found, and the majority of both bipolar and stellate cells had a very dim S100β staining. Therefore, the faint S100β expression in bipolar cells triggered by conditioned media from phagocytic microglia suggested that they could be immature astrocytes.

**Figure 5.**
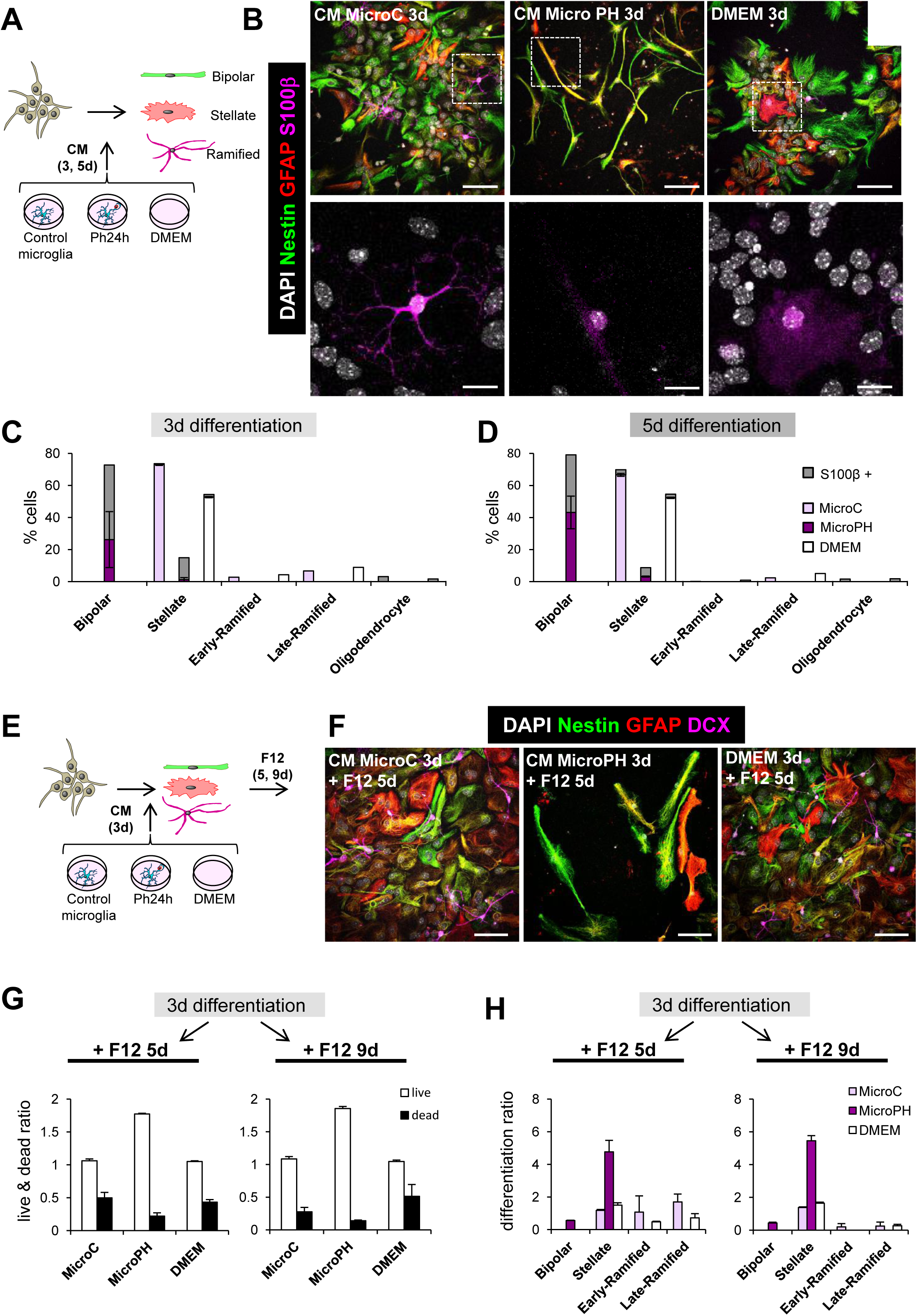
Characterization of CM cell types by S100β and multipotency assays. **[A]** Experimental design of the in vitro neurogenesis assay for S100β staining. **[B]** Representative confocal microscopy images of NPCs treated with CM microC or microPH. DMEM was used as control. **[C,D]** Percentage of expression of S100β in the different cell types found after 3d (C) and 5d (D) treatment with CM microC or microPH. **[E]** Experimental design of the in vitro multipotency assay. **[F]** Representative confocal microscopy images of NPCs treated with CM microC, microPH or DMEM followed by 5d of DMEM/F12. **[G]** Ratio of live/dead cell density over the cells at 3d after each treatment. **[H]** Differentiation ratio of each phenotype after 3d treatment with CM microC or microPH followed by 5d or 9d DMEM/F12. Scale bars, 20µm (B, F; inserts in B, 10μm); z=9 µm (B, F)N=2 independent experiments (C, D, G, H). Bars represent mean ± SEM.

Second, we analyzed the multipotency of bipolar cells. After 3d of differentiation into stellate, ramified, and bipolar cells in CM microC and microPH, we switched the culture media to DMEM/F12 culture media (the regular media to grow neurospheres) without trophic factors and allowed cells to differentiate for another 5-9d (Figure 5E-H). We found that after 5-9 days in DMEM/F12, cells derived from microPH CM had lower rates of cell death than cultures derived from microC and DMEM treatments, calculated as a ratio over the number of cells at 3 days in each culture (Figure 5G). We then analyzed the multipotency of the cultures by calculating the ratio of change of each cell type after DMEM/F12. CM microC-derived cultures presented similar ratios of stellate cells and ramified cells after 5-9d in DMEM/F12 (Figure 5H). In contrast, cultures derived from microPH CM had a strong increase in the ratio of differentiation into stellate cells while neuroblasts were not found (Figure 5H). These data strongly suggest that bipolar cells produced after treatment with the phagocytic microglia secretome were unlikely to be prototypical neuroprogenitors since they only gave rise to astrocytes but not to neuron-committed cells.

Third, we examined the calcium response of bipolar, stellate, and freshly dissociated NPCs to different stimuli using Fura-2 AM, a cell permeant calcium indicator: KCl, which triggers an intracellular Ca^+2^ response in excitable cells (De Melo Reis et al., 2011); AMPA (α-amino-3-hydroxy-5-methyl-4-isoxazolepropionic acid), which depolarizes neurons expressing the corresponding glutamate receptors (Bloodgood and Sabatini, 2008); ATP (adenosine triphosphate), which activates purinergic receptors in astrocytes and neurons (De Melo Reis et al., 2011); histamine, which triggers intracellular Ca^+2^ response in immature cells through histamine receptor, highly expressed on immature/stem cells and embryonic stem cells (Eiriz et al., 2011); and NMDA (N-methyl-D-aspartate) (Figure 6A-D). We found that 69% of the freshly dissociated NPCs depolarized in response to ATP and histamine, and became hyperpolarized in response to KCl. The majority of stellate cells depolarized in response to KCl, AMPA and ATP, and the majority of ramified cells responded to all stimuli except NMDA, most likely because they were still immature neuroblasts. On the other hand, in CM microPH treated cultures, the majority of bipolar cells highly depolarized when incubated with ATP and hyperpolarized when incubated with KCl (Figure 6C, D). These data show that bipolar cells have similar features as both NPCs and astrocytes, suggesting an intermediate phenotype.

**Figure 6.**
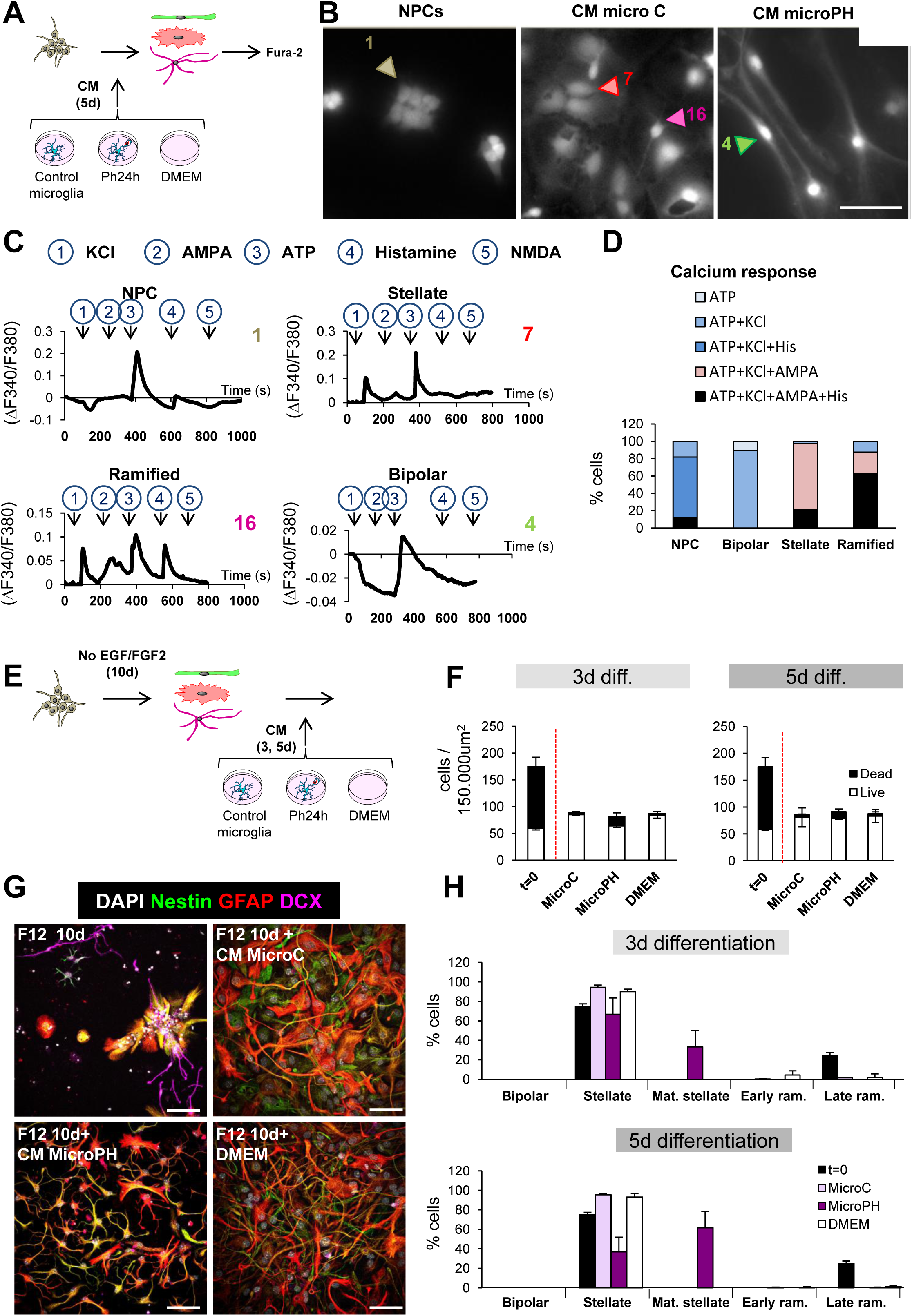
Characterization of CM cell types by calcium imaging and late survival/differentiation assays. **[A]** Experimental design of the in vitro calcium imaging assay. NPCs were treated with CM microC or microPH for 5d and the resulting stellate, ramified and bipolar cells were incubated and loaded with Fura-2 AM and afterwards, cells were challenged with KCl, AMPA, ATP, histamine and NMDA in order to measure their Ca^+2^ response. **[B]** Representative epifluorescence microscopy images of neuroprogenitors treated with CM microC or microPH for 5d. Freshly dissociated NPCs were used as control. **[C]** Calcium responses to consecutive stimuli (KCl, AMPA, ATP, Histamine, NMDA) determined as a ratio of Fura2 fluorescence of cells shown in B. **[D]** Percentage of cell phenotypes responding to each stimulus (38 stellate cells, 8 ramified cells, 19 bipolar cells, and 33 NPCs pooled from N=2 independent experiments). The baseline was calculated as the mean of the first 60sec of recording for each cell. Only peaks that increase or decrease 3 times the SEM of the baseline were considered as a positive response. **[E]** Experimental design of the in vitro late survival and differentiation assay. **[F]** Density of live and dead cells found after 10d of DMEM/F12 followed by 3-5d of CM microC or microPH and DMEM. The number of cells before adding the CM (t=0) is shown as a control. **[G]** Representative confocal microscopy images of NPCs treated with 10d of DMEM/F12 followed by 3-5d of CM microC, microPH, or DMEM. The upper left image represents the DMEM/F12 treatment of 10d, prior to adding any CM. **[H]** Percentage of cell types found after 10d of DMEM/F12 followed by 3-5d of CM microC or microPH and DMEM. The number of cells before adding the CM (t=0) is shown as a control. Mat. stellate designs stellate cells with mature (more branched) morphology, and early/late ram. designs early/late ramified cells. Scale bars, 20µm (B, E); z= 6.3µm (B, E). N=2 independent experiments (C, F, G). Bars represent mean ± SEM. ** indicates p < 0.01; *** indicates p < 0.001 by Holm-Sidak posthoc test (after one-way ANOVA was significant at p < 0.05).

Finally, we performed Western Blot analysis of NPCs as well as CM microC and CM microPH cultures of different fate-committing transcription factors: REST (RE1 silencing transcription factor 1), a repressor of neuronal genes that is highly expressed in astrocytes (Kohyama et al., 2010); Ascl (ASC1 like protein), which is related to neuronal fate commitment (Liu et al., 2015); and SMAD1 which is highly phosphorylated in differentiating astrocytes (Kohyama et al., 2010). In the CM-treated NPC cultures, we found no differences between for REST and Ascl, but the pSMAD/SMAD ratio was significantly increased in CM microPH treated cells compared to CM microC **(Supplementary Figure 5A, B)**, suggesting that CM microPH bipolar cells were committed to the astrocytic lineage. All together, these in vitro data demonstrate that at the cellular level, the microglial phagocytosis secretome promotes the inhibition of neural-committed cells which resulted in the differentiation of NPCs into astrocytes.

### Cytokines are unlikely to drive the bipolar phenotype triggered by phagocytic microglia conditioned media

We next focused on characterizing the nature of the phagocytosis secretome. As observed in the arrays (Figure 3), phagocytic microglia expressed the mRNA of several cytokines, such as Csf3, IL-1β, IL-6, TNF-α, or TGF-β among others. In addition to phagocytosis, these molecules are also released by microglia upon inflammatory stimuli and some have already been reported to impair neurogenesis (Ekdahl et al., 2003; Monje et al., 2003). To directly compare cytokine expression induced by phagocytosis and by a classical inflammatory stimulus such as LPS (bacterial lypopolysaccharides) we treated NPCs with the CM of primary microglia that had been pre-treated with LPS (1µg/ml) (Monje et al., 2003) for 6h to trigger the inflammatory response, and then changed to fresh media for another 18h to ensure that the CM would not contain any leftover LPS. Moreover, we also added another experimental group in which we treated NPCs with the CM from apoptotic SH-SY5Y in order to discard the possibility that bipolar cells could be the result of the molecules released by apoptotic cells **(Supplementary Figure 6A)**. We found that NPC cultures treated with LPS or CM microLPS produced a majority of stellate cells and a small proportion of ramified cells, while there were no differences between the two treatments in terms of cell type proportion and numbers. In contrast, apoptotic SH-SY5Y CM treated NPCs did not differ compared to CM microC **(Supplementary Figure 6B, C)**. Neither CM microLPS nor apoptotic SH-SY5Y CM treatments gave rise to bipolar cells, strongly suggesting that cytokines were unrelated to the effect of the phagocytosis secretome on NPCs.

Unexpectedly, neither LPS nor CM microLPS reduced the number of neuroblasts compared to CM microC **(Supplementary Figure 6B, C)**. These were surprising results because pro-inflammatory cytokines are well-documented to exert detrimental consequences for neurogenesis (Ekdahl et al., 2003; Monje et al., 2003). To disregard that this discrepancy resulted from different LPS concentration or exposure time compared to prior publications, we performed a series of LPS-based experiments in which we used first, a lower LPS dosage (150ng/ml for 18h) (Fraser et al., 2010) that produced a very similar cytokine expression as phagocytosis in microglia **(Supplementary Figure 6D-F)**; second, the exact LPS dosage and time (1µg/ml, 24h) described by Monje (Monje et al., 2003), where they found a reduction in DCX^+^ cells after CM microLPS treatment **(Supplementary Figure 6G, H)**; and third, the paradigm described by Monje, who used the BV2 cell line instead of primary microglia **(Supplementary Figure 6I, J)**, although they used hippocampal NPC cultures derived from adult rats. None of the LPS or CM LPS treatments gave rise to bipolar cells, strongly suggesting that cytokines are highly unlikely to drive the bipolar phenotype triggered by the phagocytic microglia secretome.

### The phagocytosis secretome reduces neuronal differentiation

We next characterized the effect of the CMs at later stages of neurogenesis using a late survival/differentiation assay in which NPCs were allowed to differentiate for 10d into neuroblasts and astrocytes using DMEM/F12 without trophic factors. At this stage (t=0), the cultures exhibited a high percentage of cell death (65.8% ± 2.4%) and the majority of the cells had a stellate morphology. These differentiated cultures were then treated for 3d and 5d with CM microC and microPH as well as DMEM for positive control (Figure 6E, F). Importantly, treatment with microPH did not result in higher levels of apoptosis than microC or DMEM (Figure 6F). Cultures treated with CM microC presented a vast majority of stellate cells and few ramified cells. In contrast, CM microPH-treated cultures showed no DCX^+^ cells, a small percentage of stellate cells and a majority of stellate cells with a more mature morphology (more complex ramifications) (Figure 6G, H). Nonetheless, the gene array data did not support a direct induction of astrogenesis. The functions ‘gliogenesis’ and ‘glial cell differentiation’ were significantly up-regulated in the ClueGo analysis (Figure 3E), but the majority of the genes found under those categories were autologous, and therefore, their overexpression would only modulate the microglial cells expressing them. As the astrocytic lineage is the default differentiation mode of neural stem cells (Bonaguidi et al., 2011; Encinas et al., 2011), these data suggest that the phagocytosis secretome inhibited neuronal differentiation, indirectly promoting astrocyte differentiation.

### Neurogenic modulatory factors secreted by phagocytic microglia in vitro alter neurogenesis in vivo

To then confirm the neurogenic modulatory role of the phagocytosis secretome on adult hippocampal neurogenesis in vivo, we injected CM microC and microPH into the hippocampus of 2m fms-EGFP mice for 6d using osmotic minipumps. After this period, BrdU was administered to track proliferating cells and mice were sacrificed 2h later (Figure 7A). Because the conditioned media would only exert a local effect, the three tissue slices most proximal to the injection site were quantified. We observed no differences in the density of total BrdU^+^ proliferative cells in CM microPH treated mice compared to CM microC treatment (Figure 7B, C). Importantly, CM microPH did not induce apoptosis in vivo (Figure 7D). In addition, there was a trend towards decreased density of proliferating BrdU^+^ rNSCs in mice treated with CM microPH compared to CM microC (p=0.0649) (Figure 7E-G). Moreover, the density of DCX^+^ neuroblasts the proportion of the different neuroblast subpopulations (AB, CD and EF), and neuroblast proliferation did not differ between CM microC and microPH (Figure 7H-J).

**Figure 7.**
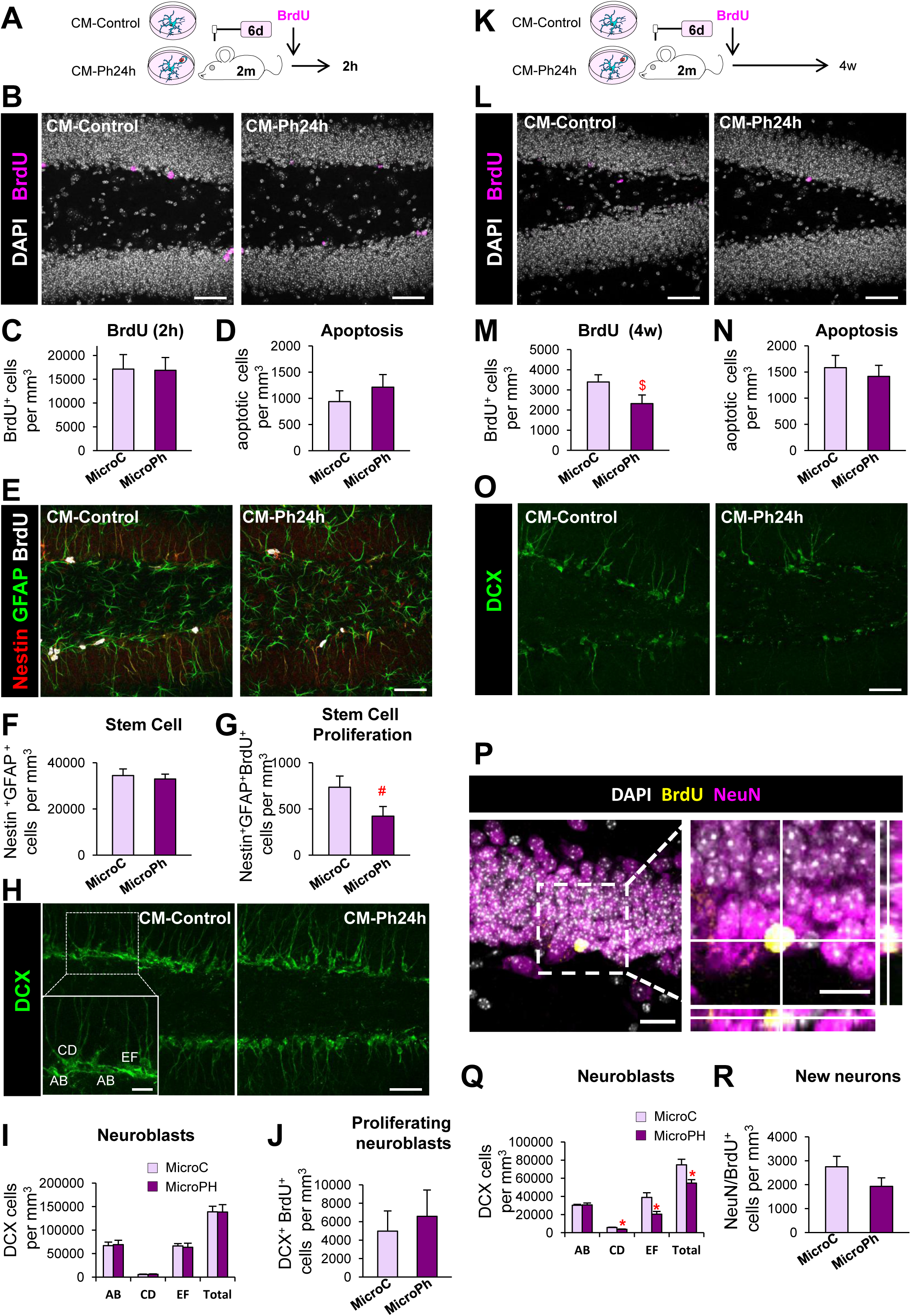
Acute and long-term effects of phagocytic microglia secreted molecules on neurogenesis in vivo. **[A]** Experimental design used for the administration of CM microC or microPH by osmotic pumps to 2m fms-EGFP mice. **[B]** Representative confocal images of cell proliferation after the CM treatments for 6d. Cell nuclei were labeled with DAPI (white) and BrdU was used as a proliferative marker (magenta). **[C]** BrdU^+^ cell density after CM microC or microPH treatment. **[D]** Apoptotic cell density after CM microC or microPH treatment. **[E]** Representative confocal images of stem cells labeled with nestin (red) and GFAP (green). **[F]** Stem cell density after CM microC or microPH treatment. **[G]** Proliferating stem cell (nestin^+^, GFAP^+^, BrdU^+^) density after CM microC or microPH treatment. **[H]** Representative confocal images of neuroblast cell populations AB, CD and EF. Neuroblast cells are labeled with DCX (green). **[I]** Density of neuroblast types AB, CD and EF. **[J]** Proliferating neuroblasts (BrdU^+^, DCX^+^) density after treatment with CM microC or microPH. **[K]** Experimental design used for the administration of CM microC or microPH by osmotic pumps to 2m fms-EGFP mice. **[L]** Representative confocal images of BrdU^+^ cells in the dentate gyrus. **[M]** BrdU^+^ cell density after CM microC or microPH treatment. **[N]** Apoptotic cell density after CM microC or microPH treatment. **[O]** Representative confocal images of neurosblasts labeled with DCX (green). **[P]** Representative confocal images of a newborn neurons labeled with BrdU (yellow) and NeuN (magenta). **[Q]** Density of neuroblast types AB, CD and EF. **[R]** New neurons (NeuN^+^, BrdU^+^) density after CM microC or microPH treatment. Scale bars, 50µm (B, L, E, O, H; insert in H, 20μm), 20µm (P; insert, 10μm); z= 12µm (B, L, E, O, H), 6μm (P). N=7-10 mice (B-I), N=3 mice (L-Q). Bars represent mean ± SEM. # indicates p=0.0649; $ indicates p=0.1257, * indicates p < 0.05; ** indicates p < 0.01; *** indicates p < 0.001 by Student’s t-test.

Because we observed a declining trend for rNSCs in the presence of CM microPH after 6d, we hypothesized that the induced alterations would accumulate over time. Therefore, we performed a long-term experiment in which mice were treated with CM microC and microPH through osmotic minipumps for 6d, followed by BrdU administration, and sacrificed 28d later to allow differentiation of the labeled cells (Figure 7K). The density of BrdU^+^ cells in mice treated with CM microPH showed a decreasing trend compared to CM microC (p=0.1257) (Figure 7L, M). Importantly, CM microPH did not induce apoptosis in vivo (Figure 7N). We then quantified neuroblasts (Figure 7O) and newborn neurons (NeuN^+^, BrdU^+^; Figure 7P). We found a significant reduction in the intermediate (CD) and most mature (EF) neuroblast subpopulations in mice treated with CM microPH compared to CM microC (Figure 7Q), while newborn neurons showed no differences in CM microPH compared to CM microC (Figure 7R; p=0.2193), possibly because few BrdU^+^ cells remained. In summary, we found a trend towards fewer proliferative stem cells at 2h after BrdU, and reduced number of mature neuroblasts and newborn neurons after 28d, suggesting that the acute (24h) phagocytic microglia secretome limits neurogenesis via the reduction in the production of neuronal-committed cells.

## DISCUSSION

In this paper, we provide evidence that microglia modulate adult hippocampal neurogenesis through the secretome associated with phagocytosis of apoptotic newborn cells, based on the following major findings. First, adult hippocampal neurogenesis was reduced in three KO models with microglial phagocytosis impairment. Second, neurogenesis was transiently increased in an acute model of phagocytosis impairment. Third, transcriptomic analysis in vitro revealed that phagocytosis triggered an expression change in a panoply of neurogenesis-related genes in microglia, strongly suggesting a coordinated neurogenic modulatory program that encompasses up to 224 heterologous genes previously shown to modulate neurogenesis, including peptides, trophic factors, matrix metalloproteases, and cytokines. Fourth, the secretome of phagocytic microglia drove NPCs differentiation towards a bipolar phenotype of astrocytic lineage in vitro, characterized by high expression of astrocytic markers such as nestin, GFAP and S100β; calcium responses to ATP and other stimuli; and high levels of phosphorylation of SMAD. Finally, the secretome of phagocytic microglia reduced the most mature neuroblast subpopulation at 28d in vivo. Hence, we present evidence that microglial phagocytosis is a pivotal mechanism for the maintenance of homeostasis in the adult neurogenic cascade.

### Microglia control the long-term homeostasis of the adult hippocampal neurogenic cascade

The accurate homeostasis of stem cell niches is crucial for their long-term maintenance, as disruption of the equilibrium between quiescence and proliferation leads to early exhaustion (Santos et al., 2018). In the adult hippocampal neurogenic niche, the mechanisms described to maintain homeostasis rely on the quiescence of rNSCs (Encinas et al., 2011), a feature that prevents an early exhaustion of the neurogenic niche (Sierra et al., 2015). Herein we focus on the unexplored role of resident immune cells, microglia, which engulf excess newborn cells that undergo apoptosis (Sierra et al., 2010). Immune cells are increasingly recognized to participate in stem cell niches, and macrophages have been recently shown to promote erythroblast production in the bone marrow (Chow et al., 2013), and to be required for ductal morphogenesis in the mammary gland (Chakrabarti et al., 2018). We found that chronic disruption of microglial phagocytosis impairs neurogenesis using two constitutive KO mice models for microglia-specific phagocytic receptors, P2Y12, and MerTK/Axl, all of which participate in different stages of phagocytosis (Elliott et al., 2009; Scott et al., 2001). Nonetheless, it is important to note that these receptors regulate multiple features of microglial physiology, and that microglial phagocytosis impairment leads to the accumulation of non-removed apoptotic cells, which may also affect neurogenesis directly through the release of toxic intracellular contents. However, the similar reduction in adult neurogenesis in the two models strongly supports the key role of microglial phagocytosis. In contrast to the effect of constitute phagocytosis impairment, acute phagocytosis impairment by inducible depletion of MerTK resulted in a transient increase in early neuroblasts that was compensated at later time points, evidencing an acute inhibitory feedback mechanism through microglial phagocytosis that is necessary for the maintenance of adult hippocampal neurogenesis in the long term.

### Microglia regulates neurogenesis through the phagocytosis secretome

We found that phagocytosis of apoptotic cells reprogrammed microglia towards a neurogenic modulatory phenotype that was mostly related to their secretome, as the majority of the modulatory genes encoded secreted proteins, including neuropeptides such as VGF, and growth factors such as VEGF and FGF2, some of which have already been described to participate in the microglial regulation of neurogenesis (Kreisel et al., 2018). In addition, the microglial secretome may contain metabolites, miRNAs and extracellular vesicles, which may also alter neurogenesis (Rodriguez-Iglesias et al., 2019). When administered in vivo, the acute secretome of phagocytic microglia inhibited hippocampal neurogenesis, as it had an early tendency to decrease rNSCs that was later followed by a reduction in mature neuroblasts. The effect was similar on isolated NPCs, as we found a decreased production of neuroblasts. However, this effect was unlikely related to an active promotion of apoptosis, which was not detected in vivo. In agreement, the in vitro transcriptomic analysis did not reveal significant increases in heterologous pro-apoptotic genes, and cell death was not observed in the late survival/differentiation assays. Nonetheless, it should be noted that many early NPCs did die upon culture with the early phagocytic microglia conditioned media, an effect that may be attributed to the lack of key survival factors, possibly metabolites consumed by phagocytic microglia. Apoptosis is common in these early cultures and has been linked to the stress associated with differentiation upon growth factor withdrawal (Babu et al., 2011). Overall, these results suggest that the secretome of phagocytic microglia modulate neurogenesis by acting not on the survival but on the differentiation of neural-committed cells.

The reduced neuronal differentiation induced by the secretome of phagocytic microglia was unlikely related to an enhancement of gliogenesis. In vivo no changes in the production of newborn astrocytes were observed in the neurogenic cascade, although on isolated NPCs the phagocytic microglial secretome gave rise to astrocyte-committed cells with a bipolar phenotype, reminiscent of radial glia (Encinas and Enikolopov, 2008). These cells presented several astrocytic features, including the expression of GFAP and S100β (Encinas and Enikolopov, 2008; Raponi et al., 2007); intracellular calcium response to ATP (De Melo Reis et al., 2011) and high phosphorylation of SMAD, which interacts with TGFβ to give rise to astrocytes/radial glia (Stipursky and Gomes, 2007). Nonetheless, our transcriptional assay did not show heterologous ‘gliogenic’ genes in phagocytic microglia, suggesting that rather than actively promoting astrogenesis, the acute phagocytosis secretome indirectly promote the default astrocytic lineage (Miller and Gauthier, 2007) by restricting the neuronal lineage.

### Cytokines are unrelated to the effect of phagocytosis secretome on neurogenesis

Several cytokines were also expressed by phagocytic microglia, such as IL-1β, IL-6, and TNFα, which have been reported to decrease survival of neuroprogenitors in vitro (IL-1β, TNFα) and in vivo (IL-6) (Breton and Mao-Draayer, 2011), inhibiting adult neurogenesis. This cytokine expression profile of phagocytic microglia holds some parallelism to the pro-inflammatory profile triggered upon inflammation, a process that has already been reported to impair neurogenesis (Ekdahl et al., 2003; Monje et al., 2003). However, in our hands the inflammatory microglia secretome did not trigger a reduction in the survival of NPCs. Furthermore, we found that both LPS and the secretome of LPS-stimulated microglia enhanced the production of neuroblasts in vitro, suggesting that inflammation is not as detrimental for neurogenesis as previously stated (Ekdahl et al., 2003; Monje et al., 2003) and that cytokines were not responsible for the effects of phagocytic microglial secretome on neural-committed cells. In addition, the neurogenic modulatory program initiated by phagocytosis encompassed genes involved in matrix remodeling (matrix metalloproteases) and membrane ligands (Jag1, ligand for Notch receptor), suggesting that the observed direct contact between microglia and rNSCs/neuroblasts (Sierra et al., 2010) may also participate in shaping the neurogenic niche through participating in the local control of neuroblast differentiation, survival and synaptic integration (Rodriguez-Iglesias et al., 2019).

### Phagocytosis reprograms microglia

Finally, we here show that phagocytosis is not simply a terminal process designed to eliminate debris. In fact, in peripheral macrophages engulfment and degradation result in epigenetic, metabolic and functional reprogramming, a process named ‘trained immunity’ (Bekkering et al., 2018). Similarly, we here show that in microglia phagocytosis of apoptotic cells triggers a coordinated transcriptional program that involves key chromatin remodeling and metabolic genes, suggesting long-term functional changes that may affect multiple microglial functions, from spine surveillance to inflammation. Apoptosis is a widespread phenomenon in neurodegenerative diseases (Abiega et al., 2016) and we speculate that phagocytosis of cell debris and the subsequent alteration of the secretome, as well as other potential functions, may be a key to understanding how microglia impacts surrounding surviving neurons. Similarly, in diseases in which microglial phagocytosis is impaired, such as epilepsy (Abiega et al., 2016), the beneficial effects of promoting engulfment/degradation of cell debris may go beyond merely removing corpses to actively promote regeneration.

In summary, in this paper, we provide strong evidence that phagocytic microglia are a central mechanism to control the homeostasis of the adult hippocampal neurogenic cascade by acutely providing a negative feedback loop via their secretome. This ‘brake’ is necessary for the long-term maintenance of the neurogenic cascade, since neurogenesis is transiently increased when phagocytosis is acutely blocked, but is disrupted when microglial phagocytosis is chronically impaired, as observed in genetically deficient mice for P2Y12 and MerTK/Axl. Importantly, the link between the proliferation of newborn cells and apoptosis was already suggested to be necessary for the correct learning and memory of the adult brain (Dupret et al., 2007), and our data here points towards microglial phagocytosis of apoptotic cells as the connecting mechanism. As apoptosis is closely related to neural stem cell proliferation, our data suggest that microglial phagocytosis may also shape other developmental and adult neurogenesis sites, such as the SVZ (Cunningham et al., 2013). In addition, phagocytosis of newborn cells has also been recently shown to play a role in sculpting sex differences in the developing amygdala (VanRyzin et al., 2019). While previous work has suggested a largely detrimental effect of microglia on hippocampal neurogenesis (Valero et al., 2016), our data are in agreement with recent evidences supporting the essential role of macrophages and other immune cells in remodeling stem cells niches (Naik et al., 2018).

## Author Contributions

IDA, IP, VST, APZ, NRI, MMR, SB, OA, LO, BS, JV and AS assisted with sample collection and sample storage/management. KH and GL provided genetic reagents and advice on their use. IDA, IP, VST, APZ, NRI, MMR, JV and AS analyzed the data. IB and IDA performed bioinformatics analysis. IDA, JV and AS, designed experiments and discussed data analysis and interpretation. IDA, IP, VST, APZ, NRI, MMR, SB, EA, CM, ASchulz, JV and AS performed experiments and analyzed results. IDA wrote the initial draft of the manuscript with input from all authors. AS and JV supervised the study and critically revised the manuscript. MMS and AS conceived the project. All authors reviewed and approved the manuscript.

## Acknowledgements

UPV/EHU SGIker technical and human support is gratefully acknowledged. We acknowledge the technical support of Victor Sánchez Zafra. We are deeply grateful to Isabel Fariñas, María Domercq and Ismael Galve-Roperh for thoughtful discussion of the data.

## SUPPLEMENTARY TEXT

Includes Methods and Methods References; Tables and Table legends; Supplementary Figures and Supplementary Figure legends

