## Supplementary Text for "Microglia actively remodels adult hippocampal neurogenesis through the phagocytosis secretome"

Includes Methods and Methods References; Tables and Table legends; Supplementary Figures and Supplementary Figure legends

### METHODS

#### CONTACT FOR REAGENT AND RESOURCE SHARING

#### EXPERIMENTAL MODEL AND SUBJECT DETAILS

**Mice:** All experiments were performed in fms-EGFP (MacGreen) mice, except where indicated, in which all microglia express the fluorescent reporter (Sasmono et al., 2003; Sierra et al., 2007). KO mice were provided by Angela Schulz (GPR34 KO), Beata Sperlagh (P2Y12 KO) and Greg Lemke (MerTK/Axl KO). Microglial-specific, inducible MerTK/Axl mice were generated using *Cx3cr1*$^{CreER}$ (Parkhurst et al., 2013) and *Mertk*$^{fl/fl}$ (Fourgeaud et al., 2016), described elsewhere. To induce deletion of the *Mertk*$^{fl/fl}$ allele in *Cx3cr1*$^{CreER/+}$*Mertk*$^{fl/fl}$ mice, two doses of tamoxifen dissolved in corn oil (75mg/kg) or corresponding volume of corn oil alone was administered i.p at p21 and p23. All mice used were in a C57BL/6 background. Mice were housed in 12:12h light:dark cycle with ad libitum access to food and water. Mice received a single dose of 5-bromo-2´-deoxyuridine (BrdU, 100-150mg/kg) at P28. At 24h or 28d after BrdU injection, mice were anesthetized with a mixture of ketamine and xylazine (100 mg/kg and 10 mg/kg, respectively), perfused with 20 U/ml heparin in PBS followed by 4% PFA in PBS. Brains were collected, immersion fixed for 4h in 4% PFA in PBS and stored in 30% sucrose, 30% ethylenglycol until analysis. All procedures followed the European Directive 2010/63/EU and NIH guidelines, and were approved by the Ethics Committees of the University of the Basque Country EHU/UPV (Leioa, Spain; CEBA/205/2011, CEBA/206/2011, CEIAB/82/2011, CEIAB/105/2012).

**SH-SY5Y cell line:** SH-SY5Y (American Type Culture Collection), a human neuroblastoma cell line derived from the bone marrow of 4 year-old female was used for phagocytic assay experiments. SH-SY5Y cells were grown as an adherent culture in non-coated culture flasks covered with 10-15ml of medium. The medium consisted of

Dulbecco's Modified Eagle Medium (DMEM, Gibco), supplemented with 10% Fetal Bovine Serum (FBS) and 1% antibiotic/antimycotic (all from Gibco). When confluency was reached, cells were trypsinized and replated at 1:4.

**BV2 cell line:** BV2 (Interlab Cell Line Collection San Martino-Instituto Scientifico Tumori-Instituto Nazionale per la Ricerca sul Cancro), a cell line derived from raf/myc-immortalized murine neonatal microglia was used to obtain LPS induced condition media. BV2 cells were grown as an adherent culture in non-coated culture flasks covered with 10-15ml of medium. The medium consisted of Dulbecco's Modified Eagle Medium (DMEM, Gibco), supplemented with 10% Fetal Bovine Serum (FBS) and 1% antibiotic/antimycotic (all from Gibco). When confluency was reached, cells were trypsinized and replated at 1:4.

**Primary Microglia Cultures:** Primary microglia cultures were performed as previously described (Abiega et al., 2016; Beccari et al., 2018). Postnatal day 0-1 (P0-P1) fms-EGFP mice pup brains were extracted and the meninges were peeled off. The olfactory bulb and cerebellum were discarded and the rest of the brain was then mechanically homogenized by careful pipetting and enzymatically digested with papain (20U/ml, Sigma), a cysteine protease enzyme, and deoxyribonuclease (DNAse; 150U/µl, Invitrogen) for 15min at 37ºC. The resulting cell suspension was then filtered through a 40µm nylon cell strainer (Fisher) and transferred to a 50ml Falcon tube quenched by 5ml of 20% Fetal Bovine Serum (FBS; Gibco) in HBSS. Afterwards, the cell suspension was centrifuged at 200g for 5min, the pellet was resuspended in 1ml Dulbecco's Modified Eagle's Medium (DMEM, Gibco) supplemented with 10% FBS and 1% Antibiotic/Antimycotic (Gibco), and seeded in T75 Poly L-Lysine-coated (15µl/ml, Sigma) culture flasks at a density of two brains per flask. Medium was changed the day after and then every 3–4 days (d), always enriched with granulocyte-macrophage colony stimulating factor (5ng/ml GM-CSF, Sigma), which promotes microglial proliferation. After confluence (at 37°C, 5% $CO_2$ for approximately 14d), microglia cells were harvested by shaking at 100-150rpm, 37°C, 4h. Isolated cells were counted and plated at a density of 80.000 cells/well on poly-l-lysine-coated glass coverslips in 24-well plates for immunofluorescence purposes or 1.000.000 cell/dish on coated Petri dishes for RT-qPCR. Microglia were allowed to settle for at least 24h before any experiment.

**NPC culture:** Neurosphere cultures were performed as previously described (Babu et al., 2011) with some modifications. Briefly, P0-P1 fms-EGFP pups were decapitated and the brains extracted and placed in cold HBSS. The homogenization process was performed as detailed in “Primary Microglia Culture Section”, except that no FBS was used in any step in order to avoid undesired neurosphere adhesion and differentiation. Afterwards, the cell suspension was centrifuged at 200g for 5min, the pellet was resuspended in 1ml DMEM/F12 with Glutamax (Gibco) supplemented with 1% Penicillin/Streptomycin, 1% B27, EGF (12,5 ng/ml), FGF-2 (5 ng/ml) (Xapelli et al., 2013). Cells were plated on uncoated Petri dishes (P60), each brain was plated in 4 Petri dishes with supplemented DMEM/F12. After six days, neurospheres were then disaggregated into a single cell suspension of NPCs using NeuroCult chemical dissociation kit following manufacturer’s instructions and each Petri dish was plated in two 6-multiwell plates. In order to maintain replicability through the experiments, neurospheres were frozen until their use at -80ºC in 15% DMSO after the first passage.

#### METHOD DETAILS

**In vitro phagocytosis assay:** The protocol is detailed here (Beccari et al., 2018). In brief, microglia were allowed to rest and settle for at least 24h before phagocytosis experiments. Phagocytosis experiments were performed in DMEM +10% FBS to ensure the presence of complement molecules, which are related to microglial phagocytosis in vivo (Diaz-Aparicio and Sierra, 2019 (under review)) and whose presence determines the immunomodulatory outcome of phagocytosis (Fraser et al., 2010). Primary microglia cells were fed for different time points with SH-SY5Y. The cell line was previously labeled with the membrane marker CM-DiI (5µM; 10min at 37°C, 15min at 4°C; Invitrogen) and treated with staurosporine (STP, 3µM, 4h; Sigma) to induce apoptosis. Only the floating dead cell fraction was collected from the supernatant and added to the primary microglia cultures in a proportion of 1:1 approximately. Apoptotic cells were visualized and quantified by trypan blue in a Neubauer chamber. Because cell membrane integrity is still maintained in early induced apoptotic cells, cells not labeled with trypan blue were considered apoptotic. The media of naïve and phagocytic (24h) microglia was immediately stored at -80ºC until its use as conditioned media for NPCs.

In some experiments, control and phagocytic microglia were treated with LPS. Three different LPS paradigms were used. In the low LPS concentration paradigm, media was removed and fresh medium with 150ng/ml LPS or vehicle (PBS) was added for 18h to primary microglia (Fraser et al., 2010). In the high LPS concentration paradigm, medium was removed and fresh medium with 1µg/ml LPS or vehicle (PBS) was added for 24h to primary or BV2 cells (Monje et al., 2003). In order to control for LPS presence in the phagocytic media, a third paradigm was performed in which primary microglia was treated with 1µg/ml LPS or vehicle (PBS) for 6h, then media was changed into fresh media for another 18h. All supernatants were collected and stored at -80ºC until its use as conditioned media for NPCs, and all of them were filter-sterilized prior to adding to the NPC culture.

**NPC proliferation and differentiation:** Neurospheres of passage 1 were thawed and expanded for a week prior to the experiment in proliferative conditions (two passages were performed in total).The day of the experiment passage 3 neurospheres were dissociated into NPCs, cells were counted and plated at a 80,000 cells/well density on poly-L-lysine coated glass coverslips in 24-well plates in supplemented (Penycillin/streptomycin, B27, EGF, and FGF2) DMEM/F12. NPCs were allowed to proliferate for 48h (Babu et al., 2011) and then were treated with conditioned media (CM) from control or phagocytic (24h) microglia. The experimental group DMEM was also added as control because it is the media in which microglia were grown. NPCs were then fixed with 4% PFA for 10min at 3d, and 5d of differentiation. For multipotency experiments, NPCs treated for 3d with conditioned media (control, 24h phagocytosis or DMEM) were transferred back to DMEM/F12 (without trophic factors) medium and were allowed to differentiate for 5d and 9d. For late survival and differentiation assay, after the 48h of proliferation, NPCs were allowed to differentiate in DMEM/F12 (no trophic factors) for 10d and then were treated with CM from control or phagocytic (24h) microglia or DMEM for another 3d and 5d.

**Calcium imaging:** Intracellular calcium imaging experiments were performed as described before (Alberdi et al., 2013). CM-treated NPCs were incubated and loaded with 5µM Fura-2 AM (Invitrogen) for 30min at 37°C and then washed in HBSS containing 20mM HEPES, pH 7.4; 10mM glucose; and 2mM $CaCl_2$ for 10min at room temperature. The perfusion chamber was assembled on the platform of a inverted epifluorescence microscope (Zeiss Axiovert 35) equipped with a 150-W xenon lamp Polychrome IV (TILL Photonics, Martinsried, Germany) and a Plan Neofluar 40× oil immersion objective (Zeiss). NPCs were treated with 50mM KCl, 10µM AMPA, 1mM ATP and 100µM histamine, sequentially. Cells were allowed to recover their baseline prior to adding the next compound. Cells were visualized with a digital black/white CCD camera (ORCA; Hamamatsu Photonics Iberica, Barcelona, Spain). Intracellular calcium signaling responses were calculated as the proportion of different cell phenotypes responding to the different stimuli. The baseline was calculated as the mean of the first 60sec of recording for each cell. Only peaks that increase or decrease 3 times the SEM of the baseline were considered as a significant response.

**FACS sorting:** Microglia cells were isolated from brains as described previously (Abiega et al., 2016; Sierra et al., 2007). The corresponding tissues from fms-EGFP mice were dissected and placed in enzymatic solution (116mM NaCl, 5.4mM KCl, 26mM $NaHCO_3$, 1mM $NaH_2PO_4$, 1.5mM $CaCl_2$, 1mM $MgSO_4$ , 0.5mM EDTA, 25mM glucose, 1mM L-cysteine) with papain (20U/ml) and DNAse I (150U/µl, Invitrogen) for digestion at 37ºC for 15min. The homogenization process was also helped by careful pipetting. After homogenization, tissue clogs were removed by filtering the cell suspension through a 40µm nylon strainer to a 50ml Falcon tube quenched by 5ml of 20% FBS in HBSS. For further enrichment of microglia, myelin was removed by using Percoll gradients. For this purpose, cells were centrifuged at 200g for 5min and resuspended in a 20% Solution of Isotonic Percoll (20% SIP; in HBSS), obtained from a previous stock of SIP (9 parts Percoll per 1 part PBS 10X). Then, each sample was layered with HBSS poured very slowly by fire-polished pipettes. Afterwards, gradients were centrifuged for 20min at 200g with minimum acceleration and no brake so the interphase was not disrupted. Then the interphase was removed, cells were washed in HBSS by centrifuging at 200g for 5min and pellet was resuspended in 500µl of sorting buffer (25mM HEPES, 5mM EDTA, 1% BSA, in HBSS). Microglia cell sorting was performed by FACS Jazz (BD), in which the population of green fluorescent cells was selected, collected in Lysis Buffer (Qiagen) containing 0.7% β-mercaptoethanol and stored at −80ºC until processing.

**Administration of microglia CM in vivo:** CM from control and phagocytic (Ph24h) microglia was administrated via osmotic pumps for 6d to 2m fms-EGFP mice. Briefly, osmotic pump (Model 2001; flow rate 1µl/h; Alzet) and infusion catheter tubes (Alzet) were filled with the conditioned media (200µl) and connected. Pumps were incubated overnight at 37ºC in PBS before the surgery. Mice were anesthetized with ketamine/xylazine (10/1 mg/kg) and received a single dose of the analgesic buprenorphine (1mg/kg) subcutaneously. The infusion cannulae were inserted at anteroposterior −1.7mm (AP), laterolateral (LL) −1.6mm, and −1.9mm dorsoventral (DV) from Bregma. Afterwards, a surface of dental cement was created from the cannulae to the screw to avoid any unwanted removal of the cannulae. Osmotic pumps were inserted inside the skin of the mice. After 6d, mice were intraperitoneally injected with BrdU (150mg/kg, single injection), and transcardically perfused 2h later to assess proliferation. For differentiation experiments, CM containing osmotic pumps were inserted for 6d to 2m fms-EGFP mice. Pumps were removed at 6d and afterwards, a single injection of BrdU (150mg/kg) was administered, and mice were sacrificed 28d later.

**Gene expression arrays:** Gene arrays analysis was performed following the recommendations of the MIAME (Minimum Information About a Microarray Experiment) consortium (Brazma et al., 2001). Cell samples from control, Ph3h and Ph24h microglia (n=3 independent experiments) were lysed and kept at -80C until processing. Total RNA was isolated using PureLink RNA Mini kit (AMBION), following manufacturer´s instructions. RNA amount was quantified in a UV/VIS NanoDrop 1000 spectrophotometer (Thermofisher), and its integrity was analyzed with Lab-chip technology in an Agilent 2100 Bioanalyzer in combination with Agilent RNA 6000 Nano Chips. Eukaryote Total RNA Nano Assay was used as type of test. In all samples, RIN>9.3, and 28S/18S>1.3 values were obtained. Sample labeling, hybridization and scanning Gene expression profiling was performed at the Gene Expression Unit of Genomics Core Facility (SGIKer) of the University of the Basque Country UPV/EHU.

One-color microarray-based gene expression analysis was performed following the One-Color (p/n5190-2305) protocol from Agilent Technologies (Low Input Quick Amp Labeling kit) for the labeling of the samples. First, 50ng of total RNA were retrotranscribed with the AffinityScript Reverse enzyme Transcriptase (AffinityScript RT), a thermostable modified enzyme derived from Moloney murine leukemia virus retrotranscriptase (Moloney Murine Leukemia Virus (MMLV) reverse transcriptase), using promoter-coupled T7 Oligo dT primers. The double-stranded cDNA synthesized by AffinityScript RT was transcribed in vitro by the T7 RNA pol in the presence of Cy3-CTP to generate labeled and amplified cRNA. The labeled samples were purified with columns of RNeasy Mini kit (Qiagen). Subsequently the labeled samples were quantified in the Nanodrop ND-1000 to determine the performance of the specific activity of the fluorochromes after labeling. All the hybridized samples met the following minimum requirements: Yield>0.825µg per reaction and Cyanine 3 specific activity>6pmol/µg.

For the hybridization, 600ng of labeled cRNA were fragmented and co-hybridized to SurePrint G3 Mouse GE 8x60K Microarray Design ID: 028005. Each array/slide contained 8 identical subarrays of more than 60.000 60-mer oligonucleotides of high resolution and performance. It contained probes for 55.681 sequences or transcripts (biological features or non-control features). Several of these biological probes were replicated 10 times for the calculations and quality control measurements (QCMetrics) of the microarrays. It also contained probes for internal positive controls (spike-ins), which were added to the RNA sample before labeling and were used for evaluation and verification of the microarray processing. Manual washing method was performed following Agilent's recommendations to prevent ozone-related problems.

Slides were scanned on a G2565CA Microarray DNA Scanner from Agilent Technologies with a resolution of 3 µm and a Tiff image size of 20bit, using the Scan software version 8.5.1 with default settings (Scan profile AgilentG3_GX_1color). The scanned TIFF images were processed and the fluorescence of the probes quantified using The Agilent Feature Extraction Software 10.7.3.1 (Agilent Technologies). Feature Extraction (FE) protocol for data extraction: GE1_107_Sep09, Design File: 28005_D_F_20140728. Software extracts information of the raw fluorescence signal (mean signal) for the fluorochrome or channel (Cy3: green Channel) from the spot containing the probes (positive and negative controls and no controls or biological feature) and the background, obtained from the negative controls (which contains sequences for which no hybridization is expected, non-specific binding indicators).

Default parameters (The Agilent Feature Extraction Software 10.7.3.1) for one-color gene expression microarrays were used for flagging of non-uniform features, population outliers for replicated probes, and features with no significant intensities in Cy3 channel. Agilent Feature Extraction (AFE) raw data was processed with software GeneSpring GX 13.0 (Agilent Technologies). Probes not present in any sample were filtered out. A list of the filtered 36.665 probes was used in the statistical analysis.

**Tissue or cultured cells RNA isolation and retrotranscription:** The corresponding tissue (P8 hippocampi for positive PCR controls) was rapidly isolated immediately under tribromoethanol overdose, and stored at -80°C. Tissue was disrupted with a roto-stator homogenizer with Lysis Buffer (Qiagen) containing 0.7% β-mercaptoethanol and stored at -80°C until processed. Cultured cells (> 500,000 cells) were lysed and stored at-80°C until processed. Total RNA was isolated using Qiagen RNeasy Mini Kit (Qiagen), following manufacturer's instructions, including a DNAse treatment step to eliminate genomic DNA residues. RNA was quantified in a Nanodrop 2000, and 1.5µg were retrotranscribed using random hexamers (Invitrogen) and Superscript III Reverse

Transcriptase kit (Invitrogen), following manufacturer's instructions in a Veriti Thermal Cycler (Applied Biosystems).

**FACS-sorted cells RNA isolation and retrotranscription:** RNA from FACS-sorted microglia (< 500,000 cells) was isolated by RNeasy Plus micro kit (Qiagen) according to the manufacturer instructions, and the RNA was retrotranscribed using an iScript Advanced cDNA Synthesis Kit (Biorad) following manufacturer instructions in a Veriti Thermal Cycler (Applied Biosystems).

**RT-qPCR:** Real Time-Quantitative Polymerase Chain Reaction (RT-qPCR) was performed following MIQE guidelines (Minimal Information for Publication of Quantitative Real Time Experiments (Bustin, 2010). Three replica of 1.5µl of a 1:3 dilution of cDNA were amplified using Power SybrGreen (Biorad) for tissue or cell culture experiments or SsoFast EvaGreen Supermix (Biorad) for FACS-sorted microglia experiments in a CFX96 Touch Real-Time PCR Detection System (Biorad). The amplification protocol for both enzymes was 3 min 95°C, and 40 cycles of 10 s at 95°C, 30 s at 60°C.

**Primers:** Primers were designed to amplify exon–exon junctions using PrimerBlast (NIH) to avoid amplification of contaminating genomic DNA, and their specificity was assessed using melting curves and electrophoresis in 2% agarose gels. Primer sequences are listed in **Supplementary Table 2**. For each set of primers, the amplification efficiency was calculated using the software LinRegPCR (Ramakers et al., 2003) or standard curve of 1:2 consecutive dilutions, and was used to calculate the relative amount using the following formula:

$$\Delta\Delta Ct = (1 + eff.target\ gene)\exp(Ct\ sample - Ct\ control)/(1 + eff.reference\ gene)\exp(Ct\ sample - Ct\ control)$$

Up to three independent reference genes were compared: L27A, which encodes a ribosomal protein of the 60S subunit (Sierra et al., 2007); OAZ-1, which encodes ornithine decarboxylase antizyme, a rate-limiting enzyme in the biosynthesis of polyamines and recently validated as reference gene in rat and human (Kwon et al., 2009); and HPRT, which encodes hypoxanthine guanine phosphoribosyl transferase (van de Moosdijk and van Amerongen, 2016). The expression of L27A, OAZ-1 and HPRT remained constant independently of time and treatments, validating their use as reference genes. In all experiments, the pattern of mRNA expression was similar using the assigned couple of reference genes, and in each experiment the reference gene that rendered lower intragroup variability was used for statistical analysis.

**Immunofluorescence**: Mice were transcardially perfused with 30ml of PBS followed by 30ml of 4% PFA. The brains were postfixed with the same fixative for 3h at RT, then washed in PBS and kept at 4°C. Six series of 50µm-thick coronal sections of mouse brains were cut using a Leica VT 1200S vibrating blade microtome (Leica Microsystems GmbH, Wetzlar, Germany). Fluorescent immunostaining was carried out following standard procedures (Beccari et al., 2018; Sierra et al., 2010). Free-floating vibratome sections were blocked in permeabilization solution (0.3% Triton-X100, 0.5% BSA in PBS; all from Sigma) for 3 hr at RT, and then incubated overnight with the primary antibodies diluted in the permeabilization solution at 4$^{o}$C. For BrdU (bromo-deoxyuridine) labeling an antigen retrieval procedure was performed by incubating in 2M HCl for 30min at 37$^{o}$C and then washing with 0.1M sodium tetraborate for 10min at RT prior to the blockade of the sections. After overnight incubation with primary antibodies (listed in **Reagents and Resources**), brain sections were thoroughly washed with 0.3% triton in PBS. Next, the sections were incubated with fluorochrome-conjugated secondary antibodies and DAPI (5mg/ml; Sigma) diluted in the permeabilization solution for 3h at RT. After washing with PBS, the sections were mounted on glass slides with DakoCytomation Fluorescent Mounting Medium (DakoCytomation, Carpinteria, CA).

Primary microglial cultures were fixed for 10min in 4% PFA and then transferred to PBS. Fluorescent immunostaining was carried out following standard procedures (Abiega et al., 2016; Beccari et al., 2018). Coverslips with primary microglial cultures were blocked in 0.1% Triton X-100, 0.5% BSA in PBS for 30min at RT. The cells were then incubated with primary antibodies in permeabilization solution (0.2% Triton X 100, 0.5% BSA in PBS) for 1h at RT, rinsed in PBS and incubated in the secondary antibodies containing DAPI (5mg/ml) in the permeabilization solution for 1h at RT. After washing with PBS, primary cultures were mounted on glass slides with DakoCytomation Fluorescent Mounting Medium (DakoCytomation, Carpinteria, CA).

For fluorouridine labeling, SH-SY5Y were treated with 2mM 5'-Fluorouridine (Sigma) for 30min. Afterwards, cells were fixed in 4% PFA with 0.5% Triton X 100. The immunofluorescence was performed as described with primary microglial cultures and anti-BrdU primary antibody was used to detect fluorouridine.

NPC cultures were fixed for 10min in 4% PFA and then transferred to PBS. Coverslips containing the cells were blocked in blocking solution (0.5% Triton-X100, 3% BSA in PBS) for 1hr at room temperature, and then incubated overnight with the primary antibodies diluted in the permeabilization solution (0.2% Triton-X100, 3% BSA in PBS) at $4^oC$. After overnight incubation, coverslips were allowed to warm at RT and were thoroughly rinsed in PBS. Next, the coverslips were incubated with fluorochrome-conjugated secondary antibodies and DAPI (5mg/ml; Sigma) diluted in the permeabilization solution for 2h at RT. After washing with PBS, the coverslips were mounted on glass slides with DakoCytomation Fluorescent Mounting Medium (DakoCytomation, Carpinteria, CA).

**Western Blot:** CM treated NPCs were directly lysed in RIPA buffer containing protease and phosphatase inhibitor cocktail (100x) (ThermoFisher). Cells were sonicated for 5s and then centrifuged (10,000g, 10min). Solubilized protein was quantified in triplicates by BCA (Bicinchoninic Acid) assay kit (ThermoFisher) at 590nm using a microplate reader (Synergy HT, BioTek). 10-15 ug of protein (denatured with β-mercaptoethanol) were loaded onto Tris-glycine gradient polyacrylamide gels (8-16%) (ThermoFisher) and run at 120V for 90min. Protein samples were then blotted to nitrocellulose membranes (0.45 um pore size) (ThermoFisher) at 220 mA for 2h. Transfer efficiency was verified by Ponceau S (Sigma) staining. For immunoblotting, membranes were rinsed in Tris Buffered Saline containing 0.1% Tween 20 (Sigma) (TBS-T) and then blocked for 1h in TBS-T containing 5% powder milk. Membranes were afterwards incubated with rabbit primary antibodies for REST (1:500, EMD Millipore), and phosphorylated Smad 1/5/9 (1:500, Cell Signaling), and mouse primary antibodies for Smad 1 (1:500, Santa Cruz), Ascl1 (1:500, BD Biosciences), and β-actin (1:5000, Sigma), in TBS-T containing 4% BSA overnight ($4^oC$, shaker). Next day, membranes were rinsed and incubated with Horseradish Peroxidase (HRP) conjugated anti-rabbit (1:5000) and anti-mouse (1:5000) secondary antibodies (Cell Signaling) in TBS-T containing 5% powder milk. After rinsing membranes, protein was visualized by enhanced chemiluminescence (ECL) using Supersignal West Femto Maximum Sensitivity Substrate (ThermoFisher) in a ChemiDoc imaging system (BioRad). Band intensity was quantified using the Gel Analyzer method of Fiji software. Phospho-Smad 1/5/9 levels were normalized to total levels of Smad 1. β-actin was used as loading control.

#### QUANTIFICATION AND STATISTICAL ANALYSIS

**Phagocytosis analysis in vivo:** The analysis of phagocytosis was performed as described (Abiega et al., 2016; Beccari et al., 2018). Apoptotic cells were defined based on their nuclear morphology after DAPI staining as cells in which the chromatin structure (euchromatin and heterochromatin) was lost and appeared condensed and/or fragmented (pyknosis/karyorrhexis). Phagocytosis was defined as the formation of an enclosed, three-dimensional pouch of microglial processes surrounding an apoptotic cell. In tissue sections, the number of apoptotic cells, phagocytosed cells, BrdU+ cells, and microglia were estimated in the volume of the DG contained in the z-stack (determined by multiplying the thickness of the stack by the area of the DG at the center of the stack using ImageJ (Fiji)). To obtain the absolute numbers, this density value was then multiplied by the volume of the septal hippocampus (spanning from −1mm to −2.5mm in the AP axes, from Bregma; approximately six slices in each of the six series), which was calculated using Fiji from a Zeiss Axiovert epifluorescent microscope images collected at 20X. For mouse tissue, 2–3 20µm-thick z-stacks containing the DG and hilus were collected per hippocampal section and a minimum of six sections per series were analyzed.

**Neurogenesis analysis in vivo:** For the neurogenesis analysis in vivo, the three tissue sections closest to the injection site were analyzed. 5 z-stacks were collected per section using a Leica SP8 laser scanning microscope under a 40X oil-immersion objective, a z-step of 1µm, a zoom of 1 and a resolution of 512x512 pixels. Proliferation was assessed by $BrdU^+$ cell quantification; neural stem cells were identified by the expression of the markers Nestin and GFAP and their radial morphology for cell quantification; neuroblast were assessed by $DCX^+$ cell quantification and morphology in order to classify them in AB, CD or EF neuroblasts (Plumpe et al., 2006); neurons were assessed by $NeuN^+$ cell quantification; and astrocytes identified as $GFAP^+$/$Nestin^-$ cells with stellate morphology. The proliferation of either of these populations was assessed by their mentioned staining combined with BrdU. Numbers of cells were estimated in the volume of the DG of the z-stack, which was determined by multiplying the thickness of the z-stack (12µm) by the area of the DG at the center of the stack using the software ImageJ (Fiji).

**Phagocytosis analysis in vitro:** Apoptotic cells were defined based on their nuclear morphology after DAPI staining as cells in which the chromatin structure (euchromatin and heterochromatin) was lost and appeared condensed and/or fragmented (pyknosis/karyorrhexis) (Abiega et al., 2016; Sierra et al., 2010). In addition, phagocytosis was defined as the formation of an enclosed, three-dimensional pouch of microglial processes surrounding an apoptotic cell. In microglia primary cultures, over 4-5 random z-stacks were collected per coverslip using a Leica SP8 laser scanning microscope under a 40X oil-immersion objective and a z-step of 0.7μm. The percentage of phagocytic microglia was defined as cells with pouches containing apoptotic SH-SY5Y nuclei and/or CM-DiI particles (Beccari et al., 2018).

**NPC differentiation analysis in vitro:** 4-5 random z-stacks were collected per coverslip using an Olympus Fluoview or a Leica SP8 laser scanning microscope under a 40X oil-immersion objective, a z-step of 0.7μm, a zoom of 0.75 and a resolution of 1024x1024 pixels. The effect of microglia-derived conditioned media on neuroprogenitor cells was analyzed considering both their morphology and the expression of cell-specific markers. Percentages of the different morphologies present in the population were obtained as well as the percentages of the different cell markers (nestin, GFAP, DCX, S100β) per morphology.

**Statistical analysis:** SigmaPlot (San Jose, CA, USA) was used for statistical analysis. Two-sample experiments were analyzed by Students´ t-Test and more than two-sample experiments with ANOVA. For the analysis of neurocandidates and cytokine mRNA expression, a logarithmic transformation was performed to comply with ANOVA assumptions (Normality and homoscedasticity). In all cases, Holm-Sidak method was used as a posthoc test to determine the significance between groups in each factor. Only $p < 0.05$ is reported to be significant. Data is shown as mean ± SEM (standard error of the mean).

**Statistical analysis of gene expression arrays**: To analyze the differential expression between naïve (t=0) and phagocytic (t=3h and t=24h) microglia groups over time the statistical analysis the maSigPro package of R/Bioconductor was used (R version 3.0.3, Bioconductor release version 2.13, maSigPro version 1.34.1) (Conesa et al., 2006). This method is based on a general regression approximation for the modeling and adjustment of the parameters required according to the type of analysis. The parameter "Time" is considered as a continuous variable, and creates a regression model of the gene response. The analysis was performed in three steps. First, genes that exhibited changes in expression over time were selected based on a p-corrected Benjamini-Hochberg (FDR) value. Next, for each of the genes that presented a significant change in their expression over time a regression was applied to determine their model (R-squared (rsq)>0.7) in order to identify patterns or models of change based on time variables, obtaining 20,800 probes. Finally, the probes were selected according to their fit to the regression model. Next, the number of genes with a very high differential pattern was reduced by applying a more restrictive criterion (rsq>0.9), obtaining 13,146. The rsq>0.7 list was used for the identification of neurogenesis related genes and the rsq>0.9 list was used for the study of transcriptional profile of phagocytic microglia.

#### DATA AND SOFTWARE AVAILABILITY

**Database for Annotation, Visualization and Integrated Discovery (DAVID):** The Database for Annotation, Visualization and Integrated Discovery (DAVID) (https://david.ncifcrf.gov/) v6.8 provides a comprehensive set of functional annotation tools to understand biological meaning behind large list of genes. DAVID was used to generate a gene-GO term enrichment analysis that identified enriched biological themes and to highlight the most relevant GO terms associated with the array gene list. The array gene list of rsq> 0.9 was used for this gene profile analysis. The analysis of each expression pattern was performed separately and only terms with an adjusted p-value (Benjamini-Hochberg) > 0.05 were considered significant.

**ClueGO:** ClueGO was used to generate protein pathways and to constitute the network of pathways based on the Gene Ontology and KEGG database (Bindea et al., 2013). ClueGO is a plugin of Cytoscape (http://www.cytoscape.org/) that visualizes the non-redundant biological terms for large clusters of genes in a functionally grouped network. A ClueGO network is created with kappa statistics and reflects the relationships between the terms based on the similarity of their associated genes (Mlecnik et al., 2018). Gene ontology (GO) analysis of mouse array data were performed with ClueGO v1.4 (Bindea et al., 2013). using the following parameters: enrichment/depletion two-sided hypergeometric statistical test; correction method: Benjamini-Hochberg; GO term range levels: 3–8; minimal number of genes for term selection: 10; minimal percentage of genes for term selection: 10%; κ-score threshold: 0.8; general term selection method: smallest p value; group method: κ; minimal number of subgroups included in a group: 3; minimal percentage of shared genes between subgroups: 50%.

#### METHODS REFERENCES


1. Abiega, O., Beccari, S., Diaz-Aparicio, I., Nadjar, A., Laye, S., Leyrolle, Q., Gomez-Nicola, D., Domercq, M., Perez-Samartin, A., Sanchez-Zafra, V., *et al.* (2016). Neuronal Hyperactivity Disturbs ATP Microgradients, Impairs Microglial Motility, and Reduces Phagocytic Receptor Expression Triggering Apoptosis/Microglial Phagocytosis Uncoupling. PLoS Biol *14*, e1002466.
2. Alberdi, E., Wyssenbach, A., Alberdi, M., Sanchez-Gomez, M.V., Cavaliere, F., Rodriguez, J.J., Verkhratsky, A., and Matute, C. (2013). Ca(2+) -dependent endoplasmic reticulum stress correlates with astrogliosis in oligomeric amyloid beta-treated astrocytes and in a model of Alzheimer's disease. Aging cell *12*, 292-302.
3. Babu, H., Claasen, J.H., Kannan, S., Runker, A.E., Palmer, T., and Kempermann, G. (2011). A protocol for isolation and enriched monolayer cultivation of neural precursor cells from mouse dentate gyrus. Front Neurosci *5*, 89.
4. Beccari, S., Diaz-Aparicio, I., and Sierra, A. (2018). Quantifying Microglial Phagocytosis of Apoptotic Cells in the Brain in Health and Disease. Curr Protoc Immunol, e49.
5. Bindea, G., Galon, J., and Mlecnik, B. (2013). CluePedia Cytoscape plugin: pathway insights using integrated experimental and in silico data. Bioinformatics *29*, 661-663.
6. Brazma, A., Hingamp, P., Quackenbush, J., Sherlock, G., Spellman, P., Stoeckert, C., Aach, J., Ansorge, W., Ball, C.A., Causton, H.C., *et al.* (2001). Minimum information about a microarray experiment (MIAME)-toward standards for microarray data. Nature genetics *29*, 365-371.
7. Bustin, S.A. (2010). Why the need for qPCR publication guidelines?--The case for MIQE. Methods *50*, 217-226.
8. Conesa, A., Nueda, M.J., Ferrer, A., and Talon, M. (2006). maSigPro: a method to identify significantly differential expression profiles in time-course microarray experiments. Bioinformatics *22*, 1096-1102.
9. Diaz-Aparicio, I., and Sierra, A. (2019 (under review)). C1q is tightly related to microglial phagocytosis in the hippocampus in physiological conditions. Science Matters.
10. Fourgeaud, L., Traves, P.G., Tufail, Y., Leal-Bailey, H., Lew, E.D., Burrola, P.G., Callaway, P., Zagorska, A., Rothlin, C.V., Nimmerjahn, A., *et al.* (2016). TAM receptors regulate multiple features of microglial physiology. Nature *532*, 240-244.
11. Fraser, D.A., Pisalyaput, K., and Tenner, A.J. (2010). C1q enhances microglial clearance of apoptotic neurons and neuronal blebs, and modulates subsequent inflammatory cytokine production. Journal of neurochemistry *112*, 733-743.
12. Kwon, M.J., Oh, E., Lee, S., Roh, M.R., Kim, S.E., Lee, Y., Choi, Y.L., In, Y.H., Park, T., Koh, S.S., *et al.* (2009). Identification of novel reference genes using multiplatform expression data and their validation for quantitative gene expression analysis. PloS one *4*, e6162.
13. Mlecnik, B., Galon, J., and Bindea, G. (2018). Comprehensive functional analysis of large lists of genes and proteins. Journal of proteomics *171*, 2-10.
14. Monje, M.L., Toda, H., and Palmer, T.D. (2003). Inflammatory blockade restores adult hippocampal neurogenesis. Science *302*, 1760-1765.
15. Parkhurst, C.N., Yang, G., Ninan, I., Savas, J.N., Yates, J.R., 3rd, Lafaille, J.J., Hempstead, B.L., Littman, D.R., and Gan, W.B. (2013). Microglia promote learning-dependent synapse formation through brain-derived neurotrophic factor. Cell *155*, 1596-1609.
16. Plumpe, T., Ehninger, D., Steiner, B., Klempin, F., Jessberger, S., Brandt, M., Romer, B., Rodriguez, G.R., Kronenberg, G., and Kempermann, G. (2006). Variability of doublecortin-associated dendrite maturation in adult hippocampal neurogenesis is independent of the regulation of precursor cell proliferation. BMC neuroscience *7*, 77.

17. Ramakers, C., Ruijter, J.M., Deprez, R.H., and Moorman, A.F. (2003). Assumption-free analysis of quantitative real-time polymerase chain reaction (PCR) data. Neuroscience letters *339*, 62-66.
18. Sasmono, R.T., Oceandy, D., Pollard, J.W., Tong, W., Pavli, P., Wainwright, B.J., Ostrowski, M.C., Himes, S.R., and Hume, D.A. (2003). A macrophage colony-stimulating factor receptor-green fluorescent protein transgene is expressed throughout the mononuclear phagocyte system of the mouse. Blood *101*, 1155-1163.
19. Sierra, A., Encinas, J.M., Deudero, J.J., Chancey, J.H., Enikolopov, G., Overstreet-Wadiche, L.S., Tsirka, S.E., and Maletic-Savatic, M. (2010). Microglia shape adult hippocampal neurogenesis through apoptosis-coupled phagocytosis. Cell Stem Cell *7*, 483-495.
20. Sierra, A., Gottfried-Blackmore, A.C., McEwen, B.S., and Bulloch, K. (2007). Microglia derived from aging mice exhibit an altered inflammatory profile. Glia *55*, 412-424.
21. van de Moosdijk, A.A., and van Amerongen, R. (2016). Identification of reliable reference genes for qRT-PCR studies of the developing mouse mammary gland. Sci Rep *6*, 35595.
22. Xapelli, S., Agasse, F., Sarda-Arroyo, L., Bernardino, L., Santos, T., Ribeiro, F.F., Valero, J., Braganca, J., Schitine, C., de Melo Reis, R.A., *et al.* (2013). Activation of type 1 cannabinoid receptor (CB1R) promotes neurogenesis in murine subventricular zone cell cultures. PloS one *8*, e63529.

#### TABLES

| Gene Symbol | FC Ph3h | FC Ph24h | Location | Effect on apoptosis |
|---|---|---|---|---|
| PRDX1 | 1,1 | 2 | autologous | anti-apoptotic |
| SIRT1 | 2 | 1,6 | autologous | anti-apoptotic |
| SMO | 1 | 3,6 | autologous | anti-apoptotic |
| SOD2 | 1,2 | 2 | autologous | anti-apoptotic |
| SPHK1 | 4,4 | 1,6 | autologous | anti-apoptotic |
| UBE2B | 1,4 | 1,5 | autologous | anti-apoptotic |
| XRCC5 | 2,1 | 3,8 | autologous | anti-apoptotic |
| PRNP | 1,8 | 1,3 | auto/hetero | anti-apoptotic |
| TGM2 | 5,5 | 4,5 | auto/hetero | anti-apoptotic |
| CNTF | 1,5 | 2,6 | heterologous | anti-apoptotic |
| FGF2 | 4,2 | 3,7 | heterologous | anti-apoptotic |
| FGF8 | 1,7 | 1,6 | heterologous | anti-apoptotic |
| VEGFA | 4,5 | 1,8 | heterologous | anti-apoptotic |
| RARG | 1,3 | 3 | autologous | pleiotropic |
| IL6 | 18,7 | 15,8 | heterologous | pleiotropic |
| BAD | 1,8 | 2,2 | autologous | pro-apoptotic |
| FAS | 1,5 | 1,5 | autologous | pro-apoptotic |
| FOXO3 | 21 | 38 | autologous | pro-apoptotic |
| GAS1 | 2,7 | 3 | autologous | pro-apoptotic |
| NLRP3 | 12 | 19 | autologous | pro-apoptotic |
| PPP2CB | 1,5 | 1,6 | autologous | pro-apoptotic |
| PTEN | 1,1 | 1,6 | autologous | pro-apoptotic |
| RHOA | 1 | 1,7 | autologous | pro-apoptotic |
| SCRIB | 1,9 | 2,4 | autologous | pro-apoptotic |
| STK3 | 1 | 1,5 | autologous | pro-apoptotic |
| TFPT | 2,4 | 3 | autologous | pro-apoptotic |
| GAL | 1,2 | 1,8 | heterologous | pro-apoptotic |
| IL1β | 8,9 | 30,7 | heterologous | pro-apoptotic |

**Table 1. Functional analysis of the phagocytic microglia transcriptome reveals changes in apoptosis.** Classification of the genes related to death cell obtained from DAVID analysis. The genes were classified according to their FC, to the effects on microglia (autologous) or on the surrounding cells (heterologous) and to the positive or negative effect on apoptosis.

| | Gene | Gene Bank | Amplicon size | Sequence 5'-3' |
|---|---|---|---|---|
| **Reference genes** | OAZ1 | NM_008753 | 51 | Fwd AGCGAGAGTTCTAGGGTTGCC<br>Rev CCCCGGACCCAGGTTACTAC |
| | L27A | BC086939 | 101 | Fwd TGTTGGAGGTGCCTGTGTTCT<br>Rev CATGCAGACAAGGAAGGATGC |
| | HPRT | NM_013556.2 | 150 | Fwd ACAGGCCAGACTTTGTTGGA<br>Rev ACTTGCGCTCATCTTAGGCT |
| **Phagocytosis receptors** | P2Y12 | NM_027571 | 88 | Fwd GCAGAACCAGGACCATGGAT<br>Rev CTGACGCACAGGGTGCTG |
| | GPR34 | NM_011823.4 | 151 | Fwd CTTCAGGAAAGCTTCAACTC<br>Rev GTAACTATCAGGAGGAGAGC |
| | MerTK | NM_008587.1 | 131 | Fwd AAGGTCCCCGTCTGTCCTAA<br>Rev GCGGGGAGGGGATTACTTTG |
| | Axl | NM_009465.4 | 86 | Fwd GTTGGTGTCTGGAGGATGGG<br>Rev TGTGTGTCCTTATGGGCTGC |
| **Peptides & hormones** | VGF | NM_001039385.1 | 74 | Fwd CACCGGCTGTCTCTGGC<br>Rev AAGGAAGCAGAAGAGGACGG |
| | Cartpt | NM_013732.7 | 106 | Fwd GCGCTATGTTGCAGATCGAAG<br>Rev GCGTCACACATGGGGACTTG |
| | FGF2 | NM_008006.2 | 113 | Fwd CGGCTGCTGGCTTCTAAGTG<br>Rev AGTGCCACATACCAACTGGAG |
| **Trophic factors** | VEGFα | NM_001025250.3 | 88 | Fwd GGCCTCCGAAACCATGAACT<br>Rev CTGGGACCACTTGGCATGG |
| | PDGFα | NM_008808.3 | 94 | Fwd TACCCCGGGAGTTGATCGAG<br>Rev TCAGCCCCTACGGAGTCTATC |
| | IGF-1 | NM_010512.4<br>NM_184052.3<br>NM_001111274.1<br>NM_001111275.1<br>NM_001111276.1 | 122 | Fwd TTACTTCAACAAGCCCACAGG<br>Rev GTGGGGCACAGTACATCTCC |
| | EGF | NM_010113.3 | 136 | Fwd GGACTGAGTTGCCCTGACTC<br>Rev CAATATGCATGCACACGCCA |
| | GDNF | NM_010275.2 | 145 | Fwd CGCTGACCAGTGACTCCAA<br>Rev TGCCGATTCCTCTCTCTTCG |
| **Matrix protein** | Mmp3 | NM_010809.2 | 88 | Fwd ACCCAGTCTACAAGTCCTCCA<br>Rev GGAGTTCCATAGAGGGACTGA |
| **Surface ligands** | Jag1 | NM_013822.5 | 119 | Fwd TTCAGGGCGATCTTGCATCA<br>Rev CACACCAGACCTTGGAGCAG |
| | Dll4 | NM_019454.3 | 113 | Fwd GGTTACACAGTGAGAAGCCAGA<br>Rev GGCAATCACACACTCGTTCC |
| **Cytokines** | Csf3 | NM_009971.1 | 70 | Fwd GCAGCCCAGATCACCCAGAAT<br>Rev TGCAGGGCCATTAGCTTCAT |
| | IL1β | NM_000576.2 | 72 | Fwd AGATGAAGTGCTCCTTCCAGG<br>Rev GGTCGGAGATTCGTAGCTGG |
| | IL6 | NM_000600.3 | 107 | Fwd GAAAGCAGCAAAGAGGCACTG<br>Rev TTCACCAGGCAAGTCTCCTCAT |
| | TNFα | NM_000594.3 | 142 | Fwd TGCACTTTGGAGTGATCGGC<br>Rev GCTTGAGGGTTTGCTACAACA |
| | TGFβ | NM_000660.5 | 112 | Fwd TCCTGGCGATACCTCAGCAA<br>Rev CAATTTCCCCTCCACGGCTC |

**Table 2. qPCR primer sequences.** List of primers used to amplify reference genes, phagocytosis receptors, peptides & hormones, trophic factors, matrix protein, surface ligands, and cytokines. The gene name, Gene Bank accession number, amplicon size, and sequence are listed.

#### SUPPLEMENTARY FIGURES LEGENDS

**Supplementary Figure 1. Apoptosis and phagocytosis in P2Y12, MerTK/Axl KO mice and iKO mice. [A]** Apoptotic cells per septal hippocampus in 1m P2Y12 KO (A1), MerTK/Axl KO (A2). **[B]** Apoptotic cells per septal hippocampus (B1), percentage of apoptotic cells engulfed (Ph index, B2) and weighted average of the percentage of microglia with phagocytic pouches (Ph capacity, B3) in P2Y12 KO mice 4w after BrdU injection (2m). **[C]** Experimental design used to isolate microglia ($GFP^+$) vs. non-microglial cells ($GFP^-$) from 1m fms-EGFP mice using flow cytometry. First, debris was excluded using the P1 gate in FSC versus SSC (left panel). Next, gates for $GFP^+$ microglia cells (P2) and $GFP^-$ non-microglial cells (P3) were defined based on the distribution of the fms-$EGFP^+$ cells in EGFP vs. FSC (right panel). **[D]** Expression of P2Y12, MerTK, and Axl in microglia ($GFP^+$) vs. non-microglial cells ($GFP^-$) by RTqPCR in FACS-sorted cells from fms-EGFP mice hippocampi. OAZ1 (Ornithine Decarboxylase Antizyme 1) was selected as a reference gene. **[E]** Apoptotic cells per septal hippocampus in MerTK iKO mice at 1d and 4w (1 and 2m (E1 and E2), respectively) after BrdU injection. N=3-4 mice (A, E1), N=4-6 mice (B, E2), N=3 independent experiments (D, each from 8 pooled hippocampi), * indicates p<0.05, ** indicates p<0.01, *** indicates p<0.001, # indicates p<0.1 by Student´s t-Test.

**Supplementary Figure 2. Transcriptional profile of phagocytic microglia. [A]** Electropherogram obtained by a Bioanalyzer comparing the RNA profile of control and phagocytic microglia as well as apoptotic SH-SY5Y (treated with 3μm STP for 24h). **[B]** Representative confocal images of $FU^+$ active transcription sites of SH-SY5Y cells treated with STP (3μm, 4h) for apoptosis induction. Nuclei were labeled with DAPI (white), cell death was detected by pyknosis/karyorrhexis (white, DAPI, arrowheads), and transcription sites were detected by FU (red). **[C]** Experimental design of the gene expression arrays. **[D]** Principal Component Analysis (PCA) of the different replica of the samples: Control microglia, Ph3h, and Ph24h. **[E]** Hierarchical Clustering (HCL) of the different replica of the samples control microglia (blue), Ph3h (brown), and Ph24h (red). **[F]** Representation of the strategy followed to screen genes from the gene array. **[G]** mRNA expression levels of the candidates selected for validation by RTqPCR in control microglia (microC), 24h phagocytic microglia (microPH) and microglia cultured with apoptotic cell conditioned media (micro+CMapo). HPRT was selected as a reference gene. Scale bar=20μm (C). N=3 independent experiments (D E, F), N=4 independent experiments (G). Bars represent mean ± SEM. * indicates p < 0.05; ** indicates p < 0.01; *** indicates p < 0.001 and # indicates p<0.06 by HolmSidak posthoc test (after one-way ANOVA was significant at $p < 0.05$). Only significant effects are shown.

**Supplementary Figure 3. Functional analysis of phagocytic microglia by DAVID and filtering strategy followed to search for phagocytic microglia expressed neurogenic modulatory genes. [A]** Functional analysis of phagocytic microglia using DAVID software. Left axis represents the fold enrichment of each biological function and right axis represents the adjusted p-value of each GO term. Key for the GO terms: GO:0014032, neural crest cell development; GO:0045664, **regulation of neuron differentiation**; GO:0050767, **regulation of neurogenesis**; GO:0019226, transmission of nerve impulse; GO:0007049, cell cycle; GO:0051301, cell division; GO:0008283, cell proliferation; GO:0000278, mitotic cell cycle; GO:0008284, positive regulation of cell proliferation; GO:0051726, regulation of cell cycle; GO:0042127, regulation of cell proliferation; GO:0048762, mesenchymal cell differentiation; GO:0045596, negative regulation of cell differentiation; GO:0045597, positive regulation of cell differentiation; GO:0006935, chemotaxis; GO:0016477, cell migration; GO:0048870, cell motility; GO:0060485, mesenchyme development; GO:0051094, positive regulation of developmental process; GO:0060284, regulation of cell development; GO:0007517, muscle organ development; GO:0035295, tube development; GO:0007507, heart development; GO:0051094, positive regulation of developmental process; GO:0060429, epithelium development; GO:0001525, angiogenesis; GO:0001568, blood vessel development; GO:0048514, blood vessel morphogenesis; GO:0042981, regulation of apoptosis; GO:0043067, regulation of programmed cell death; GO:0006955, immune response; GO:0002520, immune system development; GO:0006954, inflammatory response; GO:0045321, leukocyte activation; GO:0002694, regulation of leukocyte activation; GO:0007626, locomotory behavior; GO:0030036, actin cytoskeleton organization; GO:0007015, actin filament organization; GO:0030029, actin filament-based process; GO:0006096, **glycolysis**; GO:0046365, monosaccharide catabolic process; GO:0006796, phosphate metabolic process; GO:0006796, phosphate metabolic process; GO:0051173, positive regulation of nitrogen compound metabolic process; GO:0045935, positive regulation of nucleobase, nucleoside, nucleotide and nucleic acid metabolic process; GO:0051254, positive regulation of RNA metabolic process; GO:0016568, **chromatin modification**; GO:0016481, negative regulation of transcription; GO:0010628, positive regulation of gene expression; GO:0045941, positive regulation of transcription; GO:0045944, positive regulation of transcription from RNA polymerase II promoter; GO:0045893, positive regulation of transcription, DNA-dependent; GO:0045449, regulation of transcription; GO:0006357, regulation of transcription from RNA polymerase II promoter; GO:0051276, **chromosome organization**; GO:0006310, DNA recombination; GO:0006260, DNA replication; GO:0006974, response to DNA damage stimulus; GO:0034660, ncRNA metabolic process; GO:0009451, RNA modification; GO:0043039, tRNA aminoacylation; GO:0006399 tRNA metabolic process; GO:0007267, cell-cell signaling; GO:0007242, intracellular signaling cascade; GO:0007243, protein kinase cascade; GO:0051056, regulation of small GTPase mediated signal transduction; GO:0007264, small GTPase mediated signal transduction; GO:0016055, Wnt receptor signaling pathway; GO:0016310, phosphorylation; GO:0043038, amino acid activation; GO:0001775, cell activation; GO:0001775, cell activation; GO:0033554, cellular response to stress; GO:0030097, hemopoiesis; GO:0044271, nitrogen compound biosynthetic process; GO:0048285, organelle fission; GO:0009891, positive regulation of biosynthetic process; GO:0010557, positive regulation of macromolecule biosynthetic process; GO:0051258, protein polymerization; GO:0050865, regulation of cell activation; GO:0044057, regulation of system process; GO:0006979, response to oxidative stress; GO:0070482, response to oxygen levels; GO:0009611, response to wounding; GO:0022613, ribonucleoprotein complex biogenesis; GO:0051225, spindle assembly; GO:0006412, translation. Left axis represents the fold enrichment of each biological function and right axis represents the adjusted p-value of each GO term. Only statistically significant changes are shown. **[B]** Diagram depicting the strategy followed to search for potential modulators of neurogenesis produced by phagocytic microglia in the arrays. The filtering started by differentiating the heterologous and autologous genes in the MANGO database. Then, GO terms related to neurogenesis were selected for the heterologous MANGO genes. Afterwards, the molecules that presented the neurogenic GO terms were searched in the array. Finally, the candidate genes were filtered only to select those that appeared extracellularly (heterologous genes), and genes with receptor activity were manually discarded.

**Supplementary Figure 4. Phagocytosis-related candidates include trophic factors and peptides and hormones. [A]** Classification of the 224 potential modulators of neurogenesis. ‘Trophic factor’ was the category with the highest percentage of genes in every regulatory pattern, however, the category ‘Peptides and Hormones’ included genes with the highest FC changes in the UP regulation pattern. **[B]** The 224 candidates classified by their identity and FC.

**Supplementary Figure 5. Characterization of CM-induced phenotypes by Western blot analysis. [A]** Representative blots showing relative levels of REST, Ascl, phospho-SMAD1/5/9, and SMAD1 in NPCs treated with CM microC or microPH for 3d. **[B]** Quantification of the relative expression of REST, Ascl and the ratio phospho-SMAD/total-SMAD in NPCs treated with CM microC or microPH for 3d. β-actin was used as a loading control. N=3 independent experiments. Bars represent mean ± SEM. ** indicates $p < 0.01$; *** indicates $p < 0.001$ by Holm-Sidak posthoc test (after one-way ANOVA was significant at $p < 0.05$).

**Supplementary Figure 6.** Effect of CM microLPS on neurogenesis in vitro. **[A]** Experimental design of the in vitro neurogenesis assay. **[B]** Representative confocal microscopy images of neuroprogenitors treated with CM MicroC, CM MicroLPS 6h + 18h (1μg/ml;) or LPS alone (1μg/ml; 24h), or CM from apoptotic SH-SY5Y cells. **[C]** Percentage of cell types found after 3d or 5d treatment with CM MicroC, CM MicroLPS (6h + 18h), LPS, or CM SH apoptotic. **[D]** mRNA expression levels of selected cytokines by RTqPCR in control microglia (microC), 24h phagocytic microglia (microPH), as well as control and phagocytic microglia treated with LPS (1□g/ml, 24h). HPRT was selected as a reference gene. **[E]** Representative confocal microscopy images of NPCs treated with CM from LPS treated microglia or LPS alone (low concentration: 150ng/ml; 18h). **[F]** Quantification of the different cell types found after 3d or 5d treatment with CM from LPS treated microglia or LPS. **[G]** Representative confocal microscopy images of NPCs treated with CM MicroLPS or LPS (1µg/ml; 24h). Scale bars, 20µm. z= 7.4µm. **[H]** Quantification of the different cell types found after 3d or 5d treatment with CM MicroLPS or LPS (high concentration: 1μg/ml; 24h). **[I]** Representative confocal microscopy images of neuroprogenitors treated with CM BV2, CM BV2 LPS high or LPS high (1μg/ml; 24h). **[J]** Quantification of the different cell types found after 3d or 5d treatment with CM BV2, CM BV2 LPS or LPS. Scale bars, 20µm (B, E, G, I); z=6.3 µm (B, E, G, I). N=4 independent experiments (C, D), N=2 independent experiments (F, H, J). Bars represent mean ± SEM. * indicates $p < 0.05$; ** indicates $p < 0.01$; and *** indicates $p < 0.001$ by Holm-Sidak posthoc test (after one-way ANOVA was significant at $p < 0.05$). Only significant effects are shown.

SF1

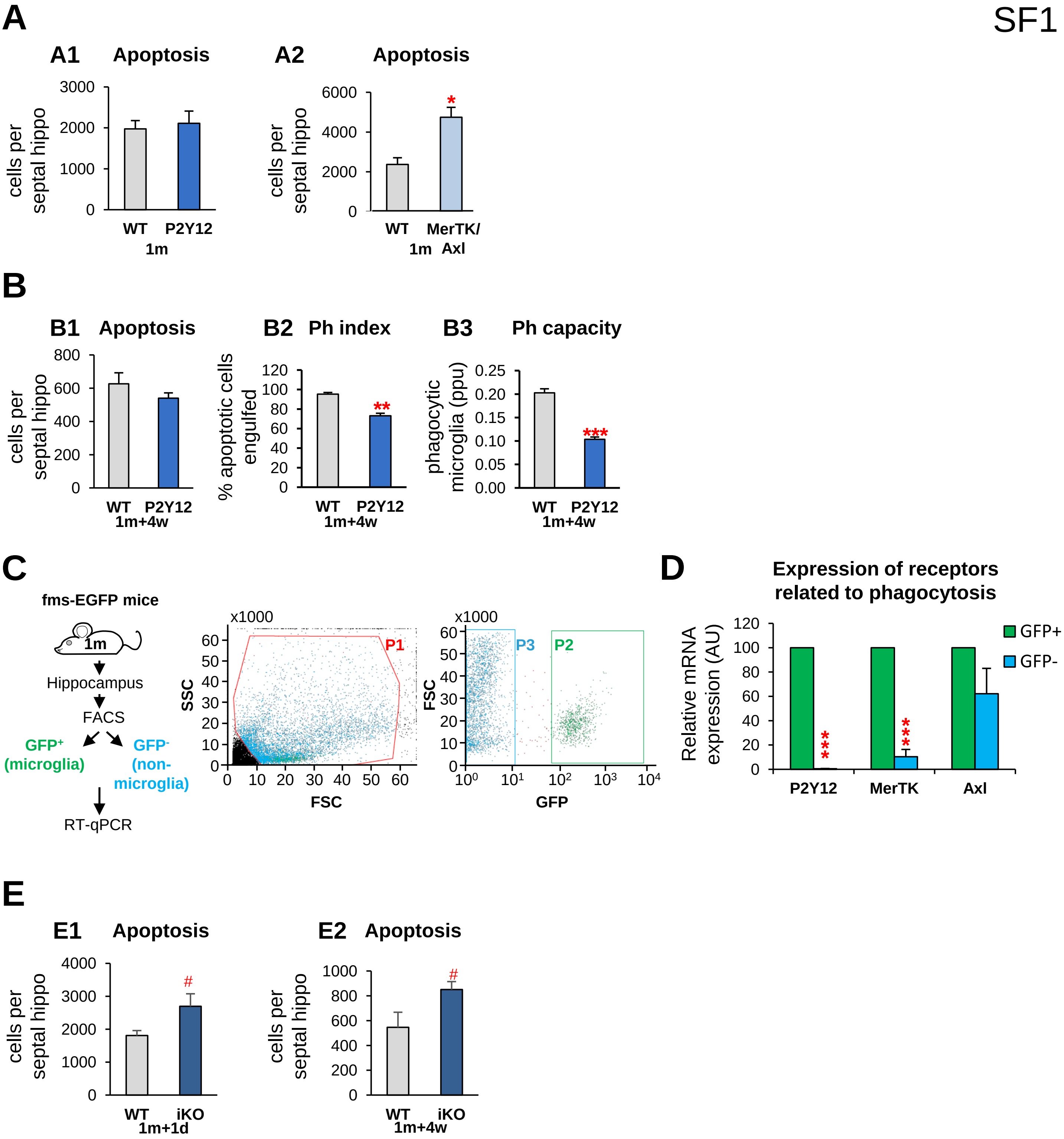
A
A1 Apoptosis
cells per septal hippo
3000
2000
1000
0
WT
P2Y12
1m
A2 Apoptosis
cells per septal hippo
6000
4000
2000
0
*
WT
MerTK/ Axl
1m
B
B1 Apoptosis
cells per septal hippo
800
600
400
200
0
WT
P2Y12
1m+4w
B2 Ph index
% apoptotic cells engulfed
120
100
80
60
40
20
0
**
WT
P2Y12
1m+4w
B3 Ph capacity
phagocytic microglia (ppu)
0.25
0.20
0.15
0.10
0.05
0.00
***
WT
P2Y12
1m+4w
C
fms-EGFP mice
1m
Hippocampus
FACS
GFP+ (microglia)
GFP- (non-microglia)
RT-qPCR
x1000
SSC
FSC
P1
x1000
FSC
GFP
P3
P2
D
Expression of receptors related to phagocytosis
Relative mRNA expression (AU)
120
100
80
60
40
20
0
GFP+
GFP-
***
***
P2Y12
MerTK
Axl
E
E1 Apoptosis
cells per septal hippo
4000
3000
2000
1000
0
#
WT
iKO
1m+1d
E2 Apoptosis
cells per septal hippo
1000
800
600
400
200
0
#
WT
iKO
1m+4w

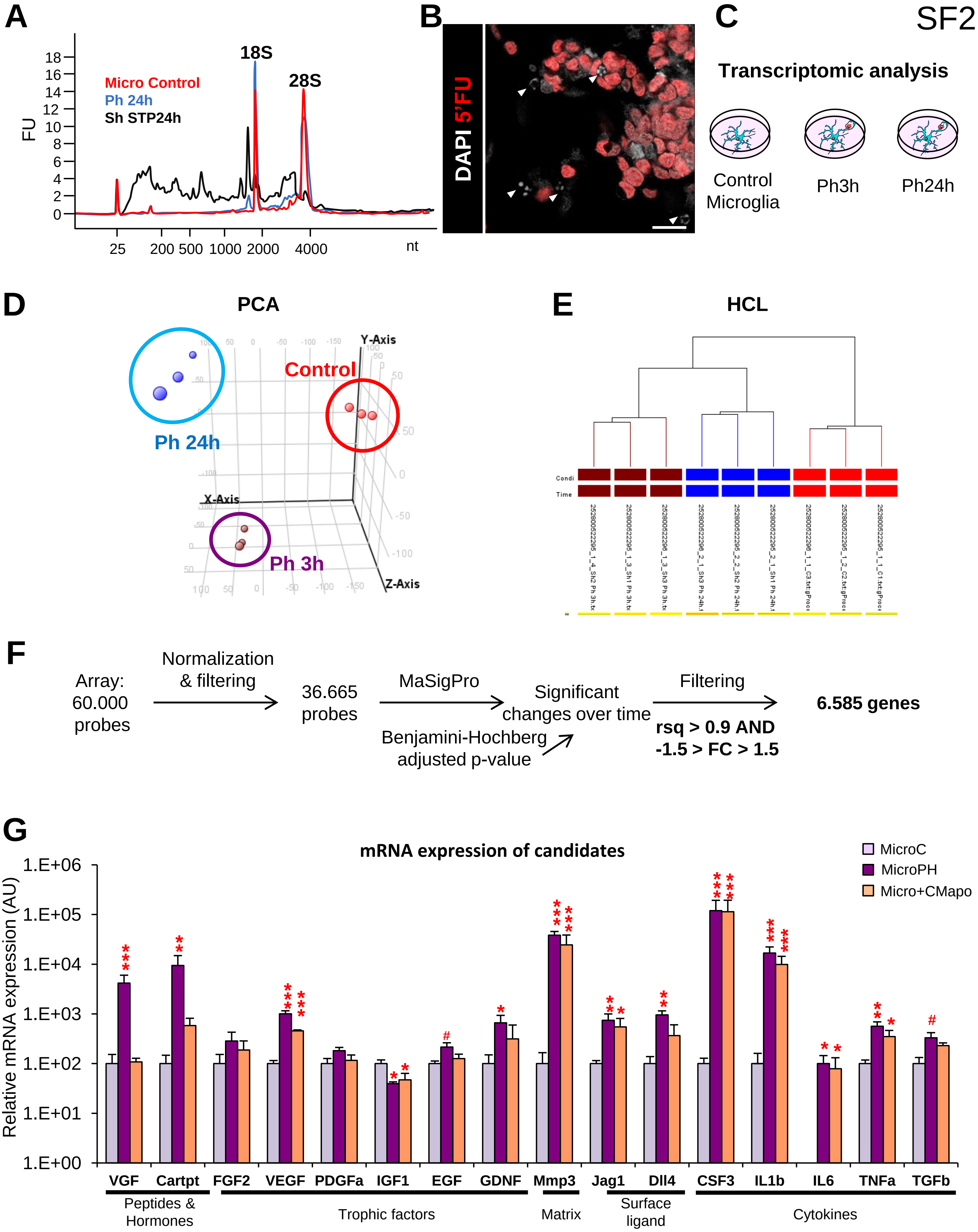
SF2
A
18S
28S
Micro Control
Ph 24h
Sh STP24h
FU
nt
B
DAPI 5'FU
C
Transcriptomic analysis
Control Microglia
Ph3h
Ph24h
D
PCA
Ph 24h
Control
Ph 3h
X-Axis
Y-Axis
Z-Axis
E
HCL
F
Array: 60.000 probes
Normalization & filtering
36.665 probes
MaSigPro
Benjamini-Hochberg adjusted p-value
Significant changes over time
Filtering
rsq > 0.9 AND -1.5 > FC > 1.5
6.585 genes
G
mRNA expression of candidates
Relative mRNA expression (AU)
MicroC
MicroPH
Micro+CMapo
VGF
Cartpt
FGF2
VEGF
PDGFa
IGF1
EGF
GDNF
Mmp3
Jag1
Dll4
CSF3
IL1b
IL6
TNFa
TGFb
Peptides & Hormones
Trophic factors
Matrix
Surface ligand
Cytokines

SF3

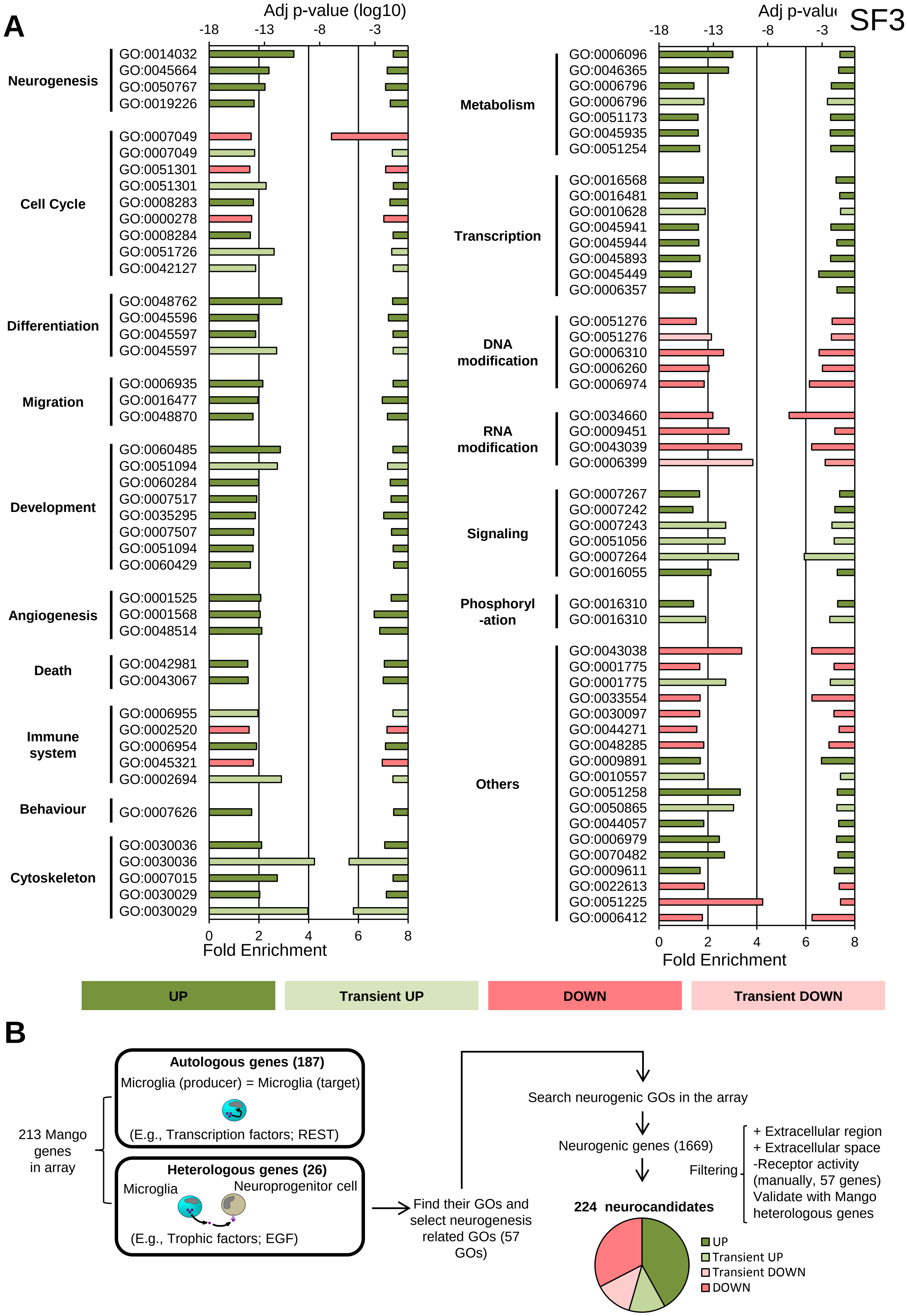
A
Adj p-value (log10)
-18
-13
-8
-3
Neurogenesis
GO:0014032
GO:0045664
GO:0050767
GO:0019226
Cell Cycle
GO:0007049
GO:0007049
GO:0051301
GO:0051301
GO:0008283
GO:0000278
GO:0008284
GO:0051726
GO:0042127
Differentiation
GO:0048762
GO:0045596
GO:0045597
GO:0045597
Migration
GO:0006935
GO:0016477
GO:0048870
Development
GO:0060485
GO:0051094
GO:0060284
GO:0007517
GO:0035295
GO:0007507
GO:0051094
GO:0060429
Angiogenesis
GO:0001525
GO:0001568
GO:0048514
Death
GO:0042981
GO:0043067
Immune system
GO:0006955
GO:0002520
GO:0006954
GO:0045321
GO:0002694
Behaviour
GO:0007626
Cytoskeleton
GO:0030036
GO:0030036
GO:0007015
GO:0030029
GO:0030029
0
2
4
6
8
Fold Enrichment
Metabolism
GO:0006096
GO:0046365
GO:0006796
GO:0006796
GO:0051173
GO:0045935
GO:0051254
Transcription
GO:0016568
GO:0016481
GO:0010628
GO:0045941
GO:0045944
GO:0045893
GO:0045449
GO:0006357
DNA modification
GO:0051276
GO:0051276
GO:0006310
GO:0006260
GO:0006974
RNA modification
GO:0034660
GO:0009451
GO:0043039
GO:0006399
Signaling
GO:0007267
GO:0007242
GO:0007243
GO:0051056
GO:0007264
GO:0016055
Phosphoryl -ation
GO:0016310
GO:0016310
Others
GO:0043038
GO:0001775
GO:0001775
GO:0033554
GO:0030097
GO:0044271
GO:0048285
GO:0009891
GO:0010557
GO:0051258
GO:0050865
GO:0044057
GO:0006979
GO:0070482
GO:0009611
GO:0022613
GO:0051225
GO:0006412
UP
Transient UP
DOWN
Transient DOWN
B
213 Mango genes in array
Autologous genes (187)
Microglia (producer) = Microglia (target)
(E.g., Transcription factors; REST)
Heterologous genes (26)
Microglia
Neuroprogenitor cell
(E.g., Trophic factors; EGF)
Find their GOs and select neurogenesis related GOs (57 GOs)
Search neurogenic GOs in the array
Neurogenic genes (1669)
Filtering
+ Extracellular region
+ Extracellular space
-Receptor activity
(manually, 57 genes)
Validate with Mango heterologous genes
224 neurocandidates
UP
Transient UP
Transient DOWN
DOWN

SF4

A

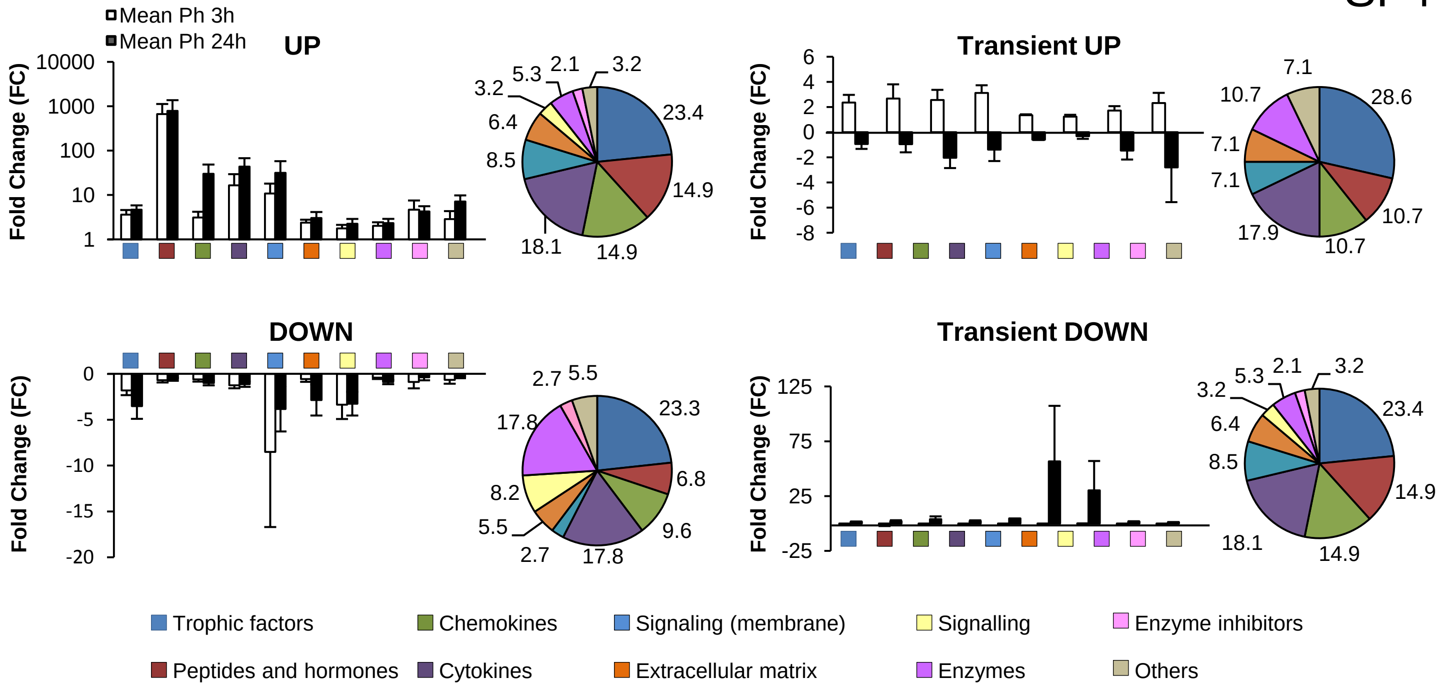


B

**Trophic factors (51genes)**

| Gene Symbol | FC Ph3h | FC Ph24h |
|---|---|---|
| Prl3c1 | 7,7 | 25,3 |
| Ereg | 5,4 | 16,0 |
| Areg | 22,6 | 5,9 |
| Inha | 1,0 | 5,6 |
| Artn | 1,0 | 5,1 |
| Bmp1 | 4,7 | 5,0 |
| Fgf23 | 3,8 | 4,4 |
| Nrtn | 2,5 | 3,7 |
| Fgf2 | 4,2 | 3,7 |
| Prl2a1 | 1,0 | 3,6 |
| Ins1 | 1,9 | 2,6 |
| Vegfa | 6,0 | 2,6 |
| Ins2 | 1,8 | 2,5 |
| Inhba | 2,3 | 2,3 |
| Prl3d1 | 1,7 | 2,3 |
| Fgf14 | 3,4 | 2,2 |
| Cntf | 1,5 | 2,2 |
| Gal | 1,2 | 1,9 |
| Fgf8 | 1,8 | 1,7 |
| Flt3l | 1,0 | 1,5 |
| Fbrs | 1,3 | 1,2 |
| Fgf18 | 1,3 | 1,2 |
| Gdf15 | 1,5 | -1,1 |
| Tgfb1 | 1,3 | -1,1 |
| Nenf | 1,1 | -1,2 |
| Hbegf | 6,1 | -1,2 |
| Pdgfa | 1,5 | -2,2 |
| Tgfb3 | 3,9 | -2,2 |
| Pdgfb | 1,7 | -2,3 |
| Cyr61 | 1,8 | -4,3 |
| Nrg4 | -1,2 | 2,2 |
| Nodal | -1,2 | 1,7 |
| Lefty2 | -1,8 | 1,4 |
| Gpi1 | -1,3 | 1,2 |
| Hdgf | -1,3 | -1,1 |
| Prl3a1 | -3,3 | -1,2 |
| Gdf9 | -2,2 | -1,3 |
| Vegfb | -1,8 | -1,3 |
| Pdgfc | -1,9 | -1,4 |
| Egfl7 | -2,6 | -1,7 |
| Grn | -1,4 | -1,7 |
| Fgf9 | -2,5 | -2,3 |
| Lefty1 | -1,2 | -2,6 |
| Bmp2 | -1,1 | -2,7 |
| Cdnf | -3,5 | -3,5 |
| Nov | -2,7 | -4,6 |
| Grem1 | -3,8 | -4,7 |
| Igf1 | -1,4 | -5,5 |
| Gdf3 | -2,5 | -7,3 |
| Vegfc | -4,7 | -9,3 |
| Ctgf | -10,0 | -24,7 |

**Peptides & Hormones (26 genes)**

| Gene Symbol | FC Ph3h | FC Ph24h |
|---|---|---|
| Vgf | 5968,2 | 7966,9 |
| Scg2 | 2931,9 | 2685,4 |
| Amh | 372,1 | 276,4 |
| Cartpt | 19,3 | 34,0 |
| Nmb | 9,9 | 19,0 |
| Edn2 | 4,0 | 7,6 |
| Npy | 1,1 | 5,3 |
| Sct | 2,3 | 3,9 |
| Retn | 4,1 | 3,8 |
| Fndc5 | 8,6 | 3,5 |
| Cort | 2,1 | 3,3 |
| Nts | 1,3 | 2,3 |
| Npff | 1,7 | 1,5 |
| Rln3 | 1,6 | 1,4 |
| Calca | 2,0 | -1,0 |
| Gnrh1 | 4,9 | -1,6 |
| Prok2 | 1,2 | -3,2 |
| Adm | -1,8 | 4,7 |
| Gcg | -1,1 | 1,5 |
| Stc2 | -5,3 | 1,5 |
| Nppa | -1,2 | 1,0 |
| Pomc | -1,8 | -1,2 |
| Insl6 | -2,3 | -1,4 |
| Edn1 | -2,2 | -1,6 |
| Hgf | -1,0 | -1,7 |
| Gnl3 | -1,2 | -2,1 |

**Chemokines (30 genes)**

| Gene Symbol | FC Ph3h | FC Ph24h |
|---|---|---|
| Cxcl3 | 6,2 | 250,1 |
| Cxcl1 | 14,5 | 104,9 |
| Cxcl2 | 8,8 | 37,5 |
| Ccl5 | 1,3 | 5,8 |
| Pf4 | 1,2 | 4,5 |
| Ccl6 | 1,1 | 3,8 |
| Ccl9 | 2,0 | 3,3 |
| Ppbp | 1,2 | 3,1 |
| Cxcl11 | 1,9 | 2,0 |
| Ccl2 | 1,1 | 1,8 |
| Ccl25 | 1,1 | 1,4 |
| Cmtm3 | 1,0 | 1,2 |
| Ybx1 | 1,3 | 1,1 |
| Ccl7 | 1,3 | 1,1 |
| Ccl12 | 2,0 | -1,4 |
| Cxcl10 | 4,1 | -3,6 |
| Cx3cl1 | 1,5 | -4,1 |
| Cxcl5 | -1,2 | 16,6 |
| Ccl3 | -3,3 | 1,8 |
| Cmtm4 | -1,7 | 1,5 |
| Cmtm7 | -1,2 | 1,5 |
| Cklf | -1,1 | 1,3 |
| Cmtm2a | -1,2 | 1,2 |
| Ccl4 | -1,4 | -1,1 |
| Cmtm8 | -1,8 | -1,3 |
| Ccl8 | -1,2 | -1,7 |
| Ccl27a | -2,1 | -1,9 |
| Cxcl14 | -1,0 | -2,2 |
| Cxcl12 | -2,7 | -2,6 |
| Cmtm5 | -1,1 | -3,1 |

**Cytokines (39 genes)**

| Gene Symbol | FC Ph3h | FC Ph24h |
|---|---|---|
| Csf3 | 10,1 | 344,2 |
| Ptn | 223,9 | 255,1 |
| Il1f6 | 2,5 | 64,2 |
| Il1b | 8,9 | 30,8 |
| Il6 | 1,2 | 18,7 |
| Il1a | 2,1 | 6,0 |
| Osm | 8,1 | 3,0 |
| Lif | 3,2 | 2,9 |
| Il19 | 2,8 | 2,7 |
| Tnfsf13b | 1,1 | 2,7 |
| Clcf1 | 3,5 | 2,2 |
| Il27 | 1,4 | 1,6 |
| Mdk | 1,2 | 1,6 |
| C3 | 1,3 | 1,6 |
| Il24 | 1,9 | 1,4 |
| Il1rn | 5,4 | 1,3 |
| Il18 | 1,6 | 1,2 |
| Ebi3 | 1,6 | -1,2 |
| Ifnb1 | 3,5 | -1,3 |
| Ltb | 3,4 | -1,7 |
| Tnfsf14 | 5,1 | -1,7 |
| Il10 | 2,1 | -6,0 |
| Il1f9 | -2,5 | 3,8 |
| Tnfsf9 | -1,7 | 2,5 |
| Nampt | -1,3 | 2,0 |
| Mif | -2,2 | 1,0 |
| Tnf | -1,9 | -1,0 |
| Il15 | -1,6 | -1,1 |
| Twsg1 | -1,2 | -1,5 |
| Aimp1 | -1,2 | -1,6 |
| Lta | -1,8 | -1,6 |
| Metrn | -1,2 | -1,8 |
| Kitl | -2,9 | -2,0 |
| Ctf1 | -3,1 | -2,0 |
| Il16 | -2,2 | -2,0 |
| Il4 | -2,4 | -2,1 |
| Tnfsf15 | -3,2 | -2,7 |
| Csf1 | -1,3 | -2,9 |
| Tnfsf10 | -5,3 | -5,2 |

**Enzymes (23 genes)**

| Gene Symbol | FC Ph3h | FC Ph24h |
|---|---|---|
| Lipg | 1,7 | 3,8 |
| Aldh3a1 | 1,1 | 3,7 |
| Enpp2 | 2,8 | 1,5 |
| Ihh | 3,1 | 1,4 |
| Furin | 1,3 | 1,3 |
| Alox5 | 2,1 | -1,6 |
| Zcchc11 | 1,0 | -1,9 |
| Plau | 2,1 | -3,9 |
| Prdx1 | -1,2 | 2,1 |
| Pla2g7 | -2,1 | 1,5 |
| Ppt1 | -1,1 | -1,0 |
| Uspl1 | -1,1 | -1,0 |
| Entpd5 | -3,2 | -1,2 |
| Arsa | -1,2 | -1,2 |
| Casp1 | -1,1 | -1,4 |
| Utp11l | -1,1 | -1,5 |
| Ctsh | -1,4 | -1,5 |
| Ang | -1,3 | -1,9 |
| Cela1 | -1,4 | -2,1 |
| Pycard | -1,2 | -2,2 |
| Ang4 | -1,4 | -2,8 |
| Htra1 | -1,2 | -3,2 |
| Ang3 | -1,4 | -3,6 |

**Signaling molecules (12 genes)**

| Gene Symbol | FC Ph3h | FC Ph24h |
|---|---|---|
| Wnt1 | 2,5 | 3,5 |
| S100a13 | 1,5 | 1,7 |
| Lgals3 | 1,3 | 1,5 |
| S100a8 | -1,0 | 83,7 |
| S100a9 | -1,2 | 6,0 |
| Hmgb2 | -1,2 | 1,0 |
| Agrp | -7,5 | -1,0 |
| Tulp3 | -1,9 | -1,3 |
| Angpt2 | -1,9 | -2,3 |
| S100b | -1,9 | -5,9 |
| Angptl3 | -2,3 | -6,4 |
| Angptl6 | -10,7 | -8,7 |

**Extracellular Matrix (15 genes)**

| Gene Symbol | FC Ph3h | FC Ph24h |
|---|---|---|
| Thbs1 | 2,8 | 8,3 |
| Lamc1 | 2,1 | 3,0 |
| Lamb2 | 1,7 | 2,6 |
| Mmp9 | 4,2 | 2,1 |
| Nog | 1,6 | 1,2 |
| Tnfaip6 | 2,0 | 1,0 |
| Adam17 | 1,4 | -1,1 |
| Sepp1 | 1,1 | -1,5 |
| Mmp3 | -1,0 | 157,9 |
| Mmp10 | -1,6 | 10,8 |
| Ecm1 | -1,1 | 1,4 |
| Lepre1 | -2,5 | -1,6 |
| Spp1 | -1,2 | -2,4 |
| Gas6 | -1,4 | -2,5 |
| Mgp | -1,2 | -8,9 |

**Membrane-bound Signaling molecules (13 genes)**

| Gene Symbol | FC Ph3h | FC Ph24h |
|---|---|---|
| Dll4 | 61,3 | 215,9 |
| Sema3b | 8,1 | 8,9 |
| Cd70 | 2,8 | 8,4 |
| Sema3f | 4,8 | 7,5 |
| Sema3a | 3,1 | 4,7 |
| Jag1 | 3,3 | 2,7 |
| Sdc1 | 1,5 | 2,0 |
| Dll3 | 1,6 | 1,7 |
| B4galt1 | 1,3 | -1,6 |
| Anxa1 | 1,4 | -1,6 |
| Sema3c | -1,4 | 4,6 |
| Sema4d | -1,3 | -2,4 |
| Efna1 | -17,7 | -7,3 |

**Others (9 genes)**

| Gene Symbol | FC Ph3h | FC Ph24h |
|---|---|---|
| Dmkn | 1,2 | 11,3 |
| Hilpda | 5,8 | 8,0 |
| Hrg | 1,6 | 2,2 |
| Pecam1 | 1,5 | -1,0 |
| Uts2b | 3,1 | -6,6 |
| Copa | -1,2 | -1,5 |
| Atxn10 | -1,2 | -1,5 |
| Aggf1 | -1,3 | -1,2 |
| Ngrn | -2,9 | -1,5 |

**Enzymes inhibitors (6 genes)**

| Gene Symbol | FC Ph3h | FC Ph24h |
|---|---|---|
| Camp | 1,8 | 5,6 |
| Serpinf2 | 7,5 | 3,0 |
| Serpine1 | -1,8 | 1,2 |
| Timp2 | -1,3 | 1,1 |
| Cst3 | -1,2 | -1,0 |
| Spink2 | -2,6 | -1,7 |

| | |
|---|---|
| FC < 2 | FC < -2 |
| FC 2-4 | FC -2-4 |
| FC > 4 | FC > -4 |

SF5

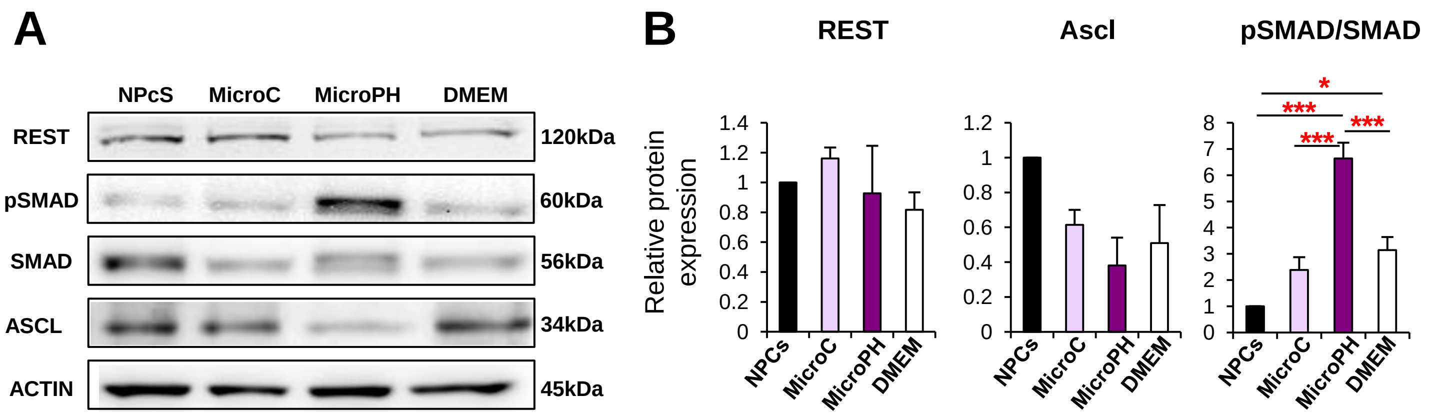

A
NPcS
MicroC
MicroPH
DMEM
REST
pSMAD
SMAD
ASCL
ACTIN
120kDa
60kDa
56kDa
34kDa
45kDa
B
REST
Ascl
pSMAD/SMAD
Relative protein expression
1.4
1.2
1
0.8
0.6
0.4
0.2
0
8
7
6
5
4
3
2
1
NPCs
MicroC
MicroPH
DMEM
*
***
***
***

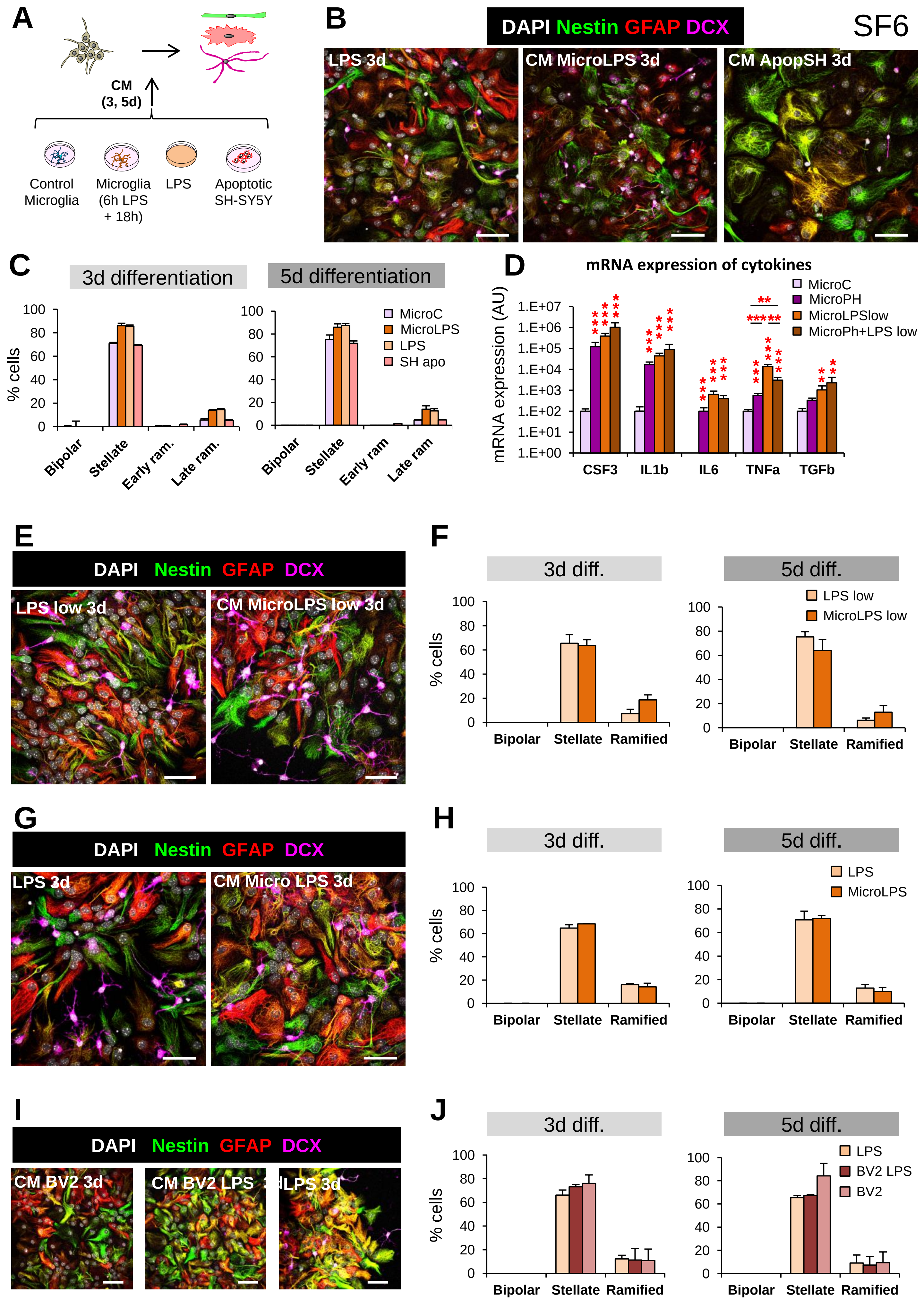

SF6
A
CM
(3, 5d)
Control Microglia
Microglia (6h LPS + 18h)
LPS
Apoptotic SH-SY5Y
B
DAPI Nestin GFAP DCX
LPS 3d
CM MicroLPS 3d
CM ApopSH 3d
C
3d differentiation
5d differentiation
% cells
MicroC
MicroLPS
LPS
SH apo
Bipolar
Stellate
Early ram.
Late ram.
D
mRNA expression of cytokines
mRNA expression (AU)
MicroC
MicroPH
MicroLPSlow
MicroPh+LPS low
CSF3
IL1b
IL6
TNFa
TGFb
E
DAPI Nestin GFAP DCX
LPS low 3d
CM MicroLPS low 3d
F
3d diff.
5d diff.
% cells
LPS low
MicroLPS low
Bipolar
Stellate
Ramified
G
DAPI Nestin GFAP DCX
LPS 3d
CM Micro LPS 3d
H
3d diff.
5d diff.
% cells
LPS
MicroLPS
Bipolar
Stellate
Ramified
I
DAPI Nestin GFAP DCX
CM BV2 3d
CM BV2 LPS 3d
LPS 3d
J
3d diff.
5d diff.
% cells
LPS
BV2 LPS
BV2
Bipolar
Stellate
Ramified
